# Mammalian epigenomic conservation of promoters and enhancers in the heart associates with trait-associated variation and impacts cardiomyocyte phenotypes

**DOI:** 10.1101/2025.06.05.658128

**Authors:** Stephanie Frost, Diego Fernandez-Aroca, Elise Parey, Adrian Rodriguez-Gonzalez, Daniel Pavon-Heredia, Diego Villar

## Abstract

Interindividual genetic variation associated with cardiovascular traits and disease is commonly found in the non-coding genome, suggesting regulatory changes underlie many genetic association signals. Mammalian gene regulation is controlled by promoters and enhancers with rapidly divergent activities. However, the interplay between human non-coding genetic variation and regulatory evolution remains largely unexplored. Here, we investigated promoter and enhancer evolution in the mammalian heart using genome-wide epigenomic profiling, intersected these elements with genetic variants associated with cardiovascular traits, and experimentally tested candidate regions in human cardiomyocytes derived from pluripotent stem cells. First, we applied a comparative genomics approach to identify epigenomically-conserved promoters and enhancers in the heart, as well as elements with primate-specific or human-only epigenomic signals. Second, we evaluated the association of common genetic variation and signatures of cell-type specific gene regulation with mammalian epigenomic conservation. We report an enrichment of common genetic variants in epigenomically-conserved promoters and enhancers, which also associate with regulatory pleiotropy across cardiac cell types, and promoter-enhancer contacts in cardiomyocytes. Based on these findings, we selected candidate epigenomically-conserved elements across three cardiovascular genetics loci (*KCNH2*, *CPEB4* and *PRKCE*), and investigated gene expression and cellular outcomes upon CRISPR/Cas9-mediated deletion of each region in human cardiomyocytes. These analyses inform how mammalian epigenomic conservation associates with specific gene expression contributions and downstream cellular responses, such as cardiomyocyte hypertrophy and sensitivity to hypoxia/reoxygenation. Moreover, the comparative analyses we present here contribute to ongoing efforts to prioritise and functionally characterise cardiovascular association signals from human population genetics.

## INTRODUCTION

Gene regulation in mammals is largely controlled by collections of non-coding regulatory elements, such as promoter and enhancer regions with characteristic biochemical hallmarks ^1,2^. Genetic alterations in specific regulatory regions are known to lead to a range of diseases, including developmental disorders and cancer ^3,4^. Moreover, human population studies over the last decade have consistently found most common disease-associated variants reside in non-coding regions of the human genome ^5–7^, further suggesting alterations in gene regulation underlie interindividual disease susceptibility ^8,9^. Whilst the contribution of selected common variants has been addressed in specific disease models ^10–12^, the recent and vast expansion of human population genetics increasingly complicates experimental investigation of individual enhancer variants.

During evolution, and in contrast to relatively stable coding sequences, mammalian regulatory elements show substantial evolutionary turnover, especially for enhancers controlling cell-type specific gene expression ^13–15^. Previous comparative epigenomic studies have extensively documented promoter and enhancer evolution across several mammalian tissues ^16–18^, and largely suggested a greater functional relevance for conserved regulatory activity. In contrast, lineage-specific elements appear to associate with low contributions to downstream gene expression, and are often compensatory to proximal losses in regulatory activity ^19^. However, only a handful of studies in mammalian tissues ^17^ or cellular models ^20^ have investigated the relationship between mammalian turnover of non-coding regulatory elements and human common genetic variation. Work across primate tissues in human and marmoset found lineage-specific enhancers were highly variable across individuals, and contributed poorly to downstream transcription ^17^. On the other hand, enhancers harbouring disease-associated variants were often conserved in human and marmoset tissues. In contrast, comparative work in human and mouse *in vitro* cardiac differentiation models ^20^ suggested disease-associated genetic variation can also occur in enhancer elements active specifically in human cardiomyocytes.

Here, we sought to characterize the evolutionary turnover of promoters and enhancers in the mammalian heart, a tissue that has been underexplored in previous comparative studies. This resource allowed us to characterize the interplay between common genetic variation associated with cardiovascular diseases and traits ^6^ in relation to epigenomic conservation of cardiac regulatory elements. Our results support previous observations on the association of trait- and disease-associated common genetic variation with conserved regulatory activity ^17^, and delineate candidate enhancers and promoters with retained epigenomic activity across mammals which also harbour human common genetic variants for cardiovascular traits. We further leveraged induced pluripotent stem cell (iPSC)-derived cardiomyocytes ^21,22^ as an *in vitro* system to investigate candidate elements via CRISPR/Cas9 genomic deletion ^23^ across three cardiovascular loci, and report the impact of each candidate element on gene expression and cellular phenotypes relevant to cardiovascular disease susceptibility, such as cardiomyocyte hypertrophy ^24^ and resistance to oxidative stress ^25^.

## RESULTS

### Heart epigenomes across 11 mammals identify a subset of epigenomically-conserved promoters and enhancers associated with core heart processes

To investigate epigenomic conservation of promoters and enhancers in the heart, we generated histone mark chromatin immunoprecipitation sequencing (ChIP-seq) data across tissue samples from ten mammalian species (Figure 1A; Figure S1). Using two to four biological replicates per species, we obtained an average of around 20,000 promoters and 50,000 enhancers reproducibly detected across individuals (Methods, Figure S1). We next used pairwise whole-genome alignments ^26^ to compare regulatory regions across our study species with those active in the human heart ^2^ (Figure 1A). First, we identified promoters and enhancers with a reciprocal alignment between the human genome and the other species’ genomes in our study phylogeny. This approach allows us to compare epigenomic signals across species in our study for the subset of alignable promoters and enhancers (Figure 1B, ∼70% of promoters and ∼60% of enhances). Second, we categorized levels of epigenomic conservation into three categories (Figure 1C). We defined epigenomically-conserved promoters and enhancers as orthologous regions displaying histone mark enrichment across all study species, which correspond to regions with highly-conserved regulatory activity. As an illustrative example, we show an epigenomically-conserved promoter and an intronic, epigenomically-conserved enhancer in the *MEF2D* cardiac locus ^27,28^ (Figure 1A). We also defined sub-sets of promoters and enhancers with lower levels of epigenomic conservation, namely restricted to primate species and to the human heart (“primate-specific” and “human-only”, respectively). In agreement with previous reports in other mammalian phylogenies and tissues ^16,29^, we identified hundreds to thousands of epigenomically-conserved and primate-specific promoters and enhancers; and larger numbers of human-only regulatory regions. Moreover, orthologous promoters with higher levels of epigenomic conservation are more abundant than enhancers in the same category (Figure 1C). Consistent with epigenomic conservation associating with core regulatory function, ontology terms across these categories showed significant enrichment of heart-related processes in epigenomically-conserved enhancers, and general metabolic and transcriptional categories for epigenomically- conserved promoters (Figure S1). These results closely agree with those reported in similar datasets ^13,16,18^, and indicate that across our study species we capture promoters and enhancers corresponding both to constrained and fast-evolving sequences.

**Figure 1:**
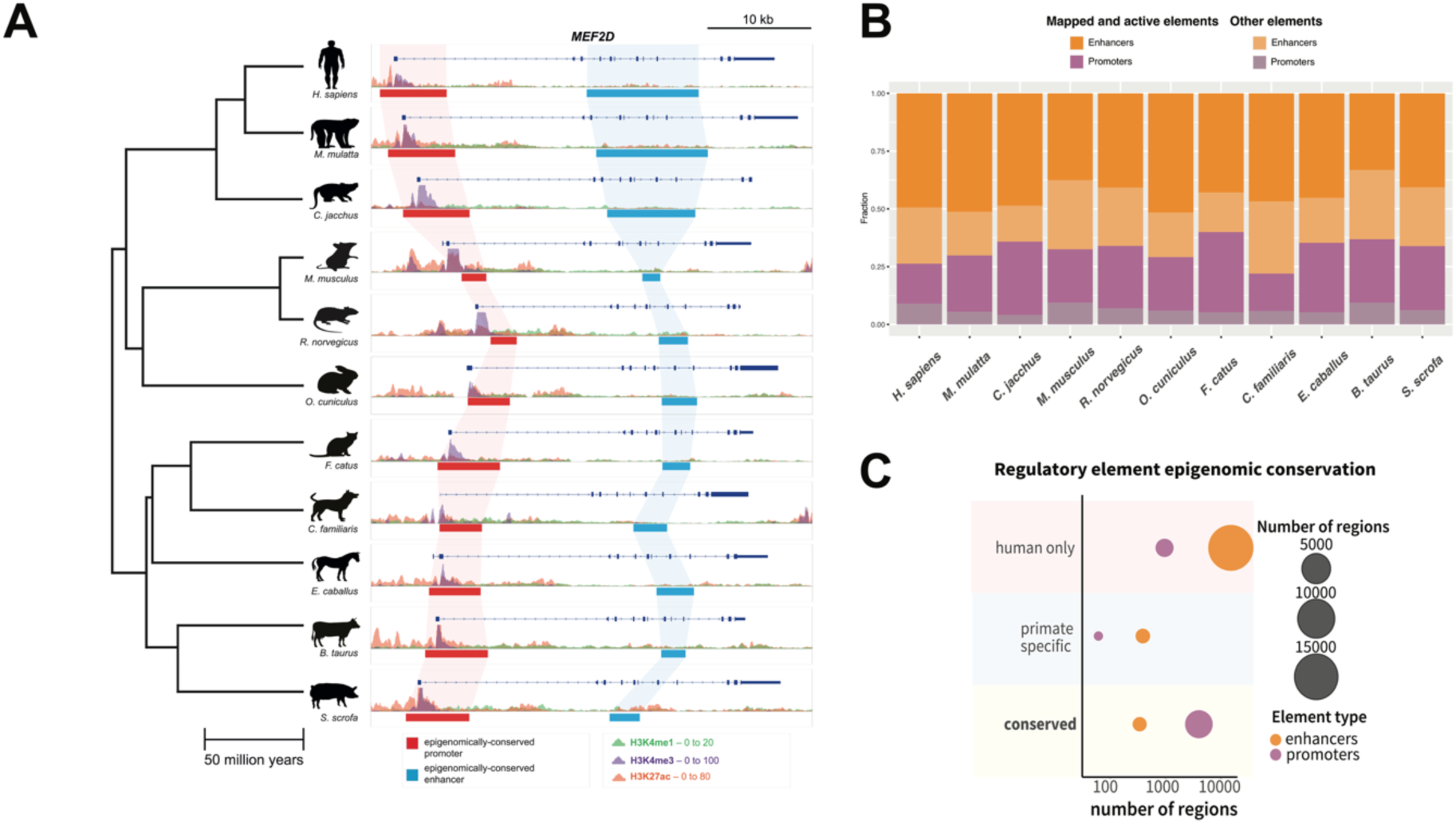
Comparative epigenomics of promoters and enhancers in the heart across in 11 mammals **A.** Epigenomic profiling and cross-mapping exemplified for the *MEF2D* cardiac locus. H3K27ac (orange), H3K4me3 (purple), and H3K4me1 (green) histone mark enrichment in heart tissue is shown for each of the species: human (*H. sapiens*), macaque (*M. mulatta*), marmoset (*C. jacchus*), mouse (*M. musculus*), rat (*R. norvegicus*), rabbit (*O. cuniculus*), cat (*F. catus*), dog (*C. familiaris*), horse (*E. caballus*), cow (*B. taurus*) and pig (*S.scrofa*). Scales next to histone marks legend indicates fold enrichment over input (averaged across replicates). Note that we use the same scale across species to facilitate direct visual comparison of histone mark enrichments. Identified promoters and enhancers showing epigenomic conservation across all species are represented by red and blue boxes, respectively. **B.** Percentage of promoters and enhancers in each species corresponding to active and mappable elements across the study species (dark purple and orange bars). Lighter shade bars indicate promoters and enhancers corresponding to non-mappable elements (Methods). **C.** Circle plot indicating numbers of promoters and enhancers across epigenomic conservation categories: epigenomically-conserved across all study species (conserved), with epigenomic conservation in primate species (primate-specific) and epigenomically- active only in human samples (human-only). See also **Figure S1** and **Table S1**.

### Epigenomic conservation in the heart enriches for trait-associated genetic variation and cell-type specific gene regulation

We next focused on characterizing the interplay between epigenomic enhancer and promoter conservation, human non-coding genetic variation and downstream gene regulation in the heart. Previous reports in primate tissues ^17^ and *in vitro* models of heart muscle ^20^ begun to explore this association, which our data addressers in an extended phylogenetic framework. To this end, we obtained common genetic variants associated with major cardiovascular traits and diseases in human population studies ^6^ (Methods), which we intersected with epigenomically-conserved, primate-specific and human-only promoters and enhancers. This analysis revealed a consistent enrichment of common genetic variation within epigenomically- conserved promoters and enhancers (Figure 2A, Table S2), with less pronounced enrichments found for primate-specific elements and cardiovascular traits such as QT interval duration. We also observed epigenomically-conserved elements associate with expression quantitative trait loci ^30^ in heart but not in liver (Table S2). Our results are consistent with previous reports in primate tissues ^17^, and suggest the association of epigenomically- conserved regulatory elements and trait-associated genetic variation can inform the contributions of heart gene regulation to cardiovascular traits.

**Figure 2:**
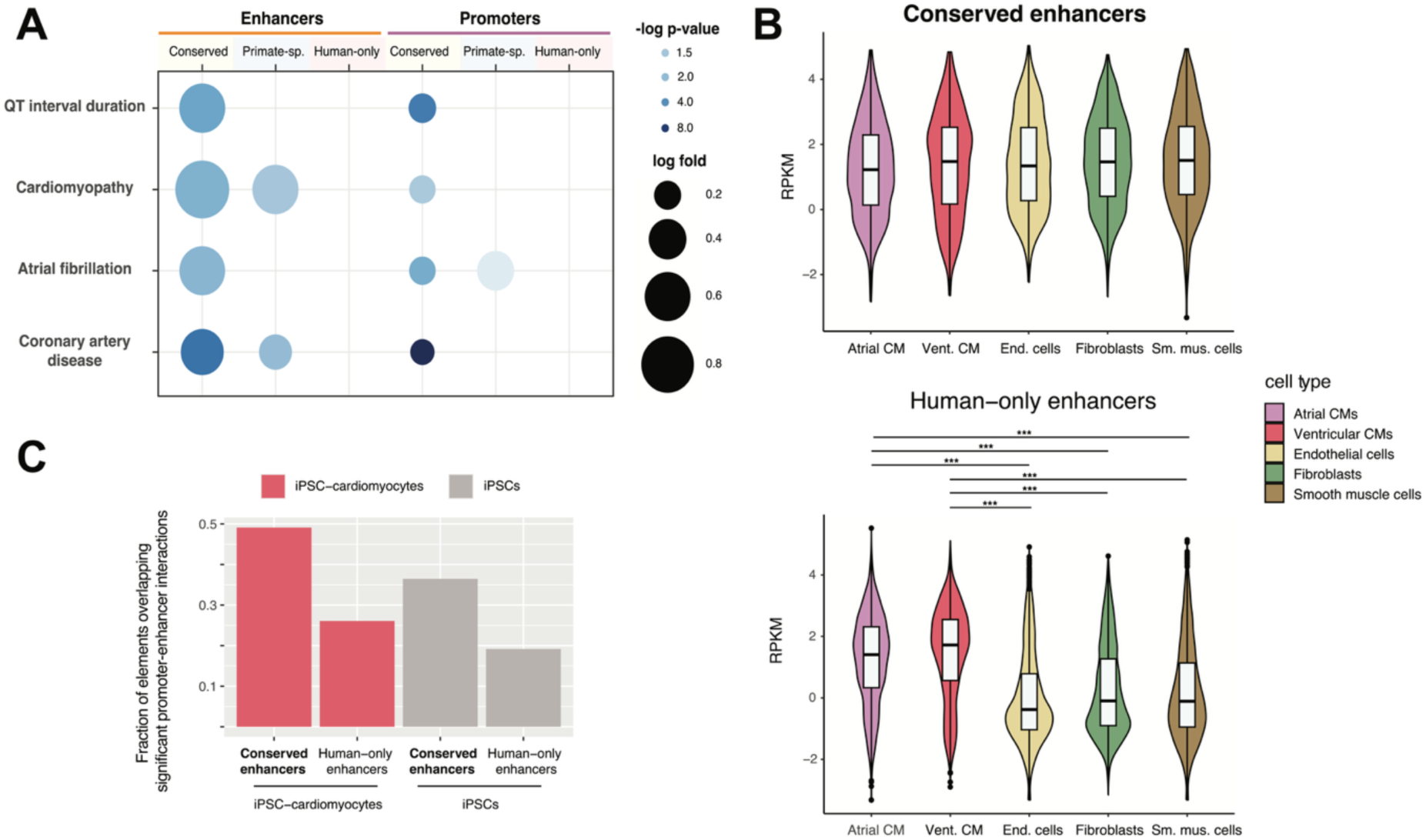
Epigenomic enhancer conservation and its association with GWAS variants and regulatory signatures in human heart cell types **A.** Enrichment of cardiovascular trait-associated common genetic variants in epigenomically- conserved and primate-specific promoters and enhancers (hypergeometric test with Benjamini–Hochberg correction for multiple testing, Methods) **B.** Chromatin accessibility across heart cell types differs between epigenomically-conserved and human-only enhancers. Mann–Whitney U tests, with Benjamini–Hochberg correction for multiple testing, (***) corrected P-values < 0.001. **C.** Promoter-enhancer contacts in iPSC-cardiomyocytes are preferentially associated with epigenomically-conserved enhancers (p = 1e-18 for iPSC-cardiomyocytes and p=1.4e-12 for iPSCs, hypergeometric test with Bonferroni correction for multiple testing, Methods). See also **Figure S2** and **Table S2**.

To further characterize the regulatory properties of epigenomically-conserved promoters and enhancers, we leveraged publicly-available datasets across human heart cell types ^31^ and iPSC-derived cardiomyocytes ^32^. First, we observed significant differences in chromatin accessibility across cardiac cell types between epigenomically-conserved and human-only elements. Whereas human-only promoters and enhancers have a higher average accessibility in cardiomyocytes, epigenomically-conserved elements show similar accessibility levels across major cardiac cell types (Figure 2B and Figure S2). This observation is consistent with earlier reports on increased regulatory pleiotropy associating with epigenomic enhancer conservation ^16,33,34^. Second, epigenomically-conserved regulatory elements are overrepresented among significant promoter-enhancer interactions in iPSC-cardiomyocytes ^32^ (p=1e-18 for enhancers and p=1e-87 for promoters; hypergeometric test with Bonferroni correction for multiple testing, Table S2). On average, 48% of epigenomically-conserved elements overlap a known promoter-enhancer interaction in this cell-type, compared with 26% of human-only enhancers (Figure 2C). On the whole, these observations support the notion that epigenomic conservation associates with downstream signatures of gene regulation across heart cell types, and in cardiomyocytes specifically.

### Genomic deletion of epigenomically-conserved enhancers and promoters during cardiomyocyte differentiation

Post-GWAS interrogation of candidate trait-associated regions from human population studies remains a significant challenge. Our previous results argue mammalian epigenomic conservation of promoters and enhancers in the heart concurs with both trait-associated genetic variation and downstream signatures of gene regulation, and may therefore delineate candidate regulatory elements for downstream perturbation experiments.

To validate this approach, we focused on candidate regions with evidence of regulatory activity in cardiomyocytes, a major cardiac cell type amenable to efficient *in vitro* differentiation from human iPSCs ^21,35^. Specifically, we employed a range of selection criteria (Methods) to define candidate genomic elements from epigenomically-conserved promoters and enhancers that harbour trait-associated genetic variation and signatures of gene regulation in human cardiomyocytes (Table S3).

From these, we selected three cardiovascular loci for targeted perturbation experiments in human induced pluripotent stem cells via CRISPR/Cas9-mediated genomic deletion (Methods, Figures 3 and S3). For each tested region, we obtained heterozygous and homozygous deletions in human iPSCs, and assessed the impact of each deletion on target gene expression at two different time-points during *in vitro* cardiomyocyte differentiation.

**Figure 3:**
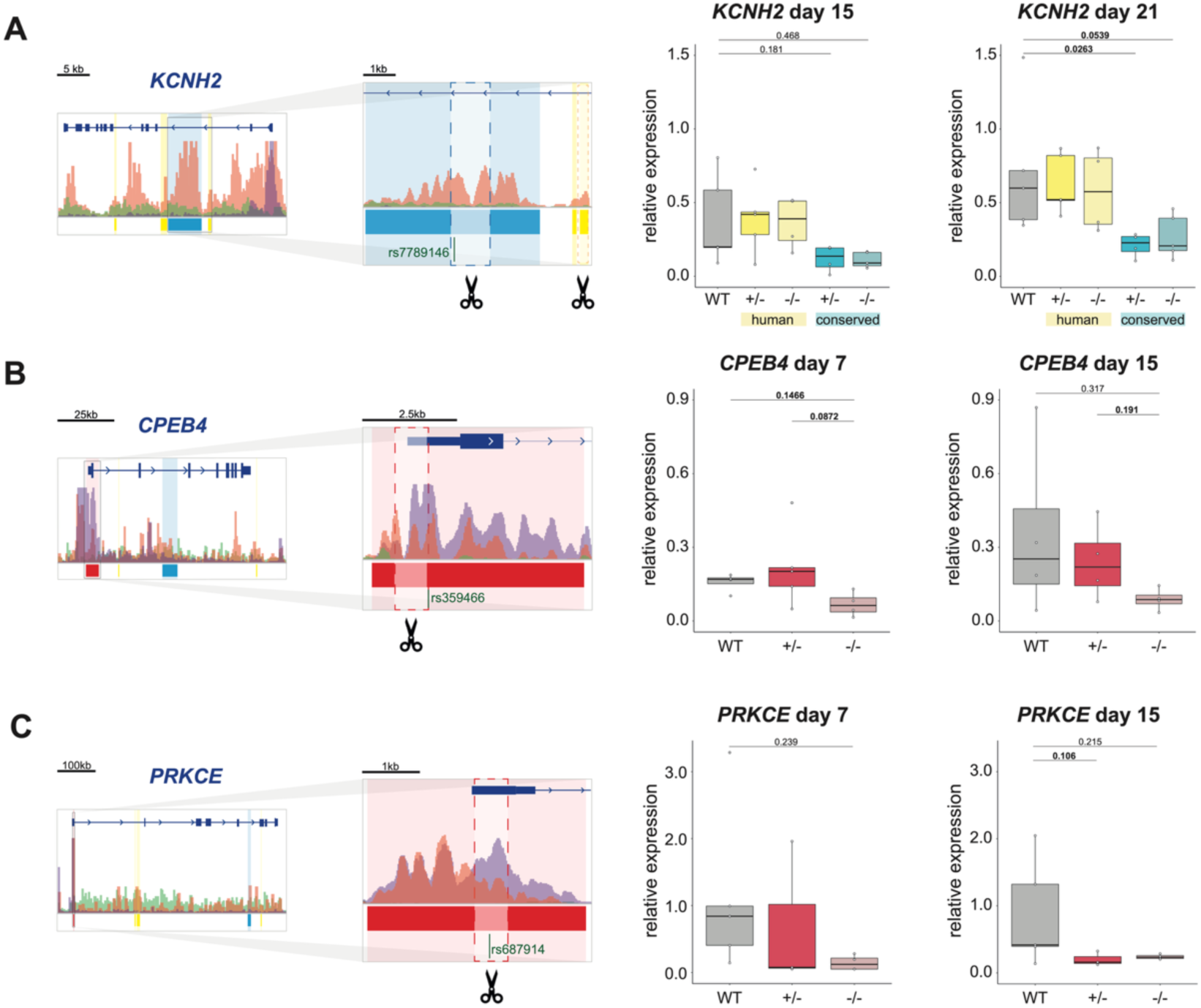
Genomic deletion of candidate promoters and enhancers across three cardiovascular loci in iPSC-cardiomyocytes **A.** Genomic deletion of an epigenomically-conserved enhancer and a control, human-only enhancer in the *KCNH2* locus. Diagrams **(left)** depict epigenomic histone mark enrichments across the gene locus, and around the two candidate enhancers (expanded inset, blue for epigenomically-conserved and yellow for human-only control). Green bars indicate selected common genetic variants associated to cardiovascular traits (Methods), in this case overlapping the first candidate enhancer. Details as in Figure 1A. Boxplots **(right)** show *KCNH2* gene expression at days 15 and 21 of cardiac differentiation, represented as relative expression over an internal reference gene (Methods) and across iPSC-cardiomyocytes with the following genotypes: wild-type (grey bar), either heterozygous (+/-) or homozygous (-/-) knock-out for epigenomically-conserved candidate enhancer (shades of blue) or the same for human-only control enhancer (shades of yellow). Highlighted pairwise p-values correspond to one-way ANOVA with Tukey post-hoc correction. **B.** Genomic deletion of an epigenomically-conserved promoter in the *CPEB4* locus (promoter 1) and assessment of changes in *CPEB4* gene expression. Details as in A. **C.** Genomic deletion of an epigenomically-conserved promoter in the *PRKCE* locus (promoter 2) and assessment of changes in *PRKCE* gene expression. Details as in A. See also **Figure S3** and **Table S3**.

First, we selected two candidate enhancers in the *KCNH2* locus: an epigenomically-conserved intronic element and a control human-only element nearby (Figure 3A). *KCNH2* codes for a transmembrane potassium channel involved in electrical conduction in cardiomyocytes ^36^, and the selected epigenomically-conserved enhancer overlaps common genetic variants associated to cardiac arrythmia ^37,38^. *KCNH2* expression at days 15 and 21 of differentiation showed negligible effects upon genomic deletion of the control, human-only enhancer (yellow, Figure 3A). In contrast, deletion of the neighbouring epigenomically-conserved enhancer led to clear reductions of *KCHN2* expression at both time-points (blue, Figure 3A). Compared to the human-only control enhancer, our results are consistent with a specific gene expression contribution of the epigenomically-conserved *KCNH2* enhancer during human cardiomyocyte differentiation, an observation that aligns with previous perturbations of overlapping or orthologous sequences in human ^31^ and mouse ^36^.

Second, we targeted an epigenomically-conserved promoter in the *CPEB4* locus. *CPEB4* codes for one of four RNA-binding proteins in the CPEB family, with reported roles in cell survival ^39^, hypoxia responses ^40^ and cardiomyocyte hypertrophy ^41^. Human population studies have found common genetic variants in this locus associated to several cardiovascular traits ^42–44^, including variants within the targeted promoter associated to myocardial mass ^7^.

Consistent with the requirement of this epigenomically-conserved promoter for gene expression, we observed a clear downregulation of *CPEB4* mRNA levels at days 7 and 15 of cardiomyocyte differentiation (Figure 3B).

Lastly, we genetically deleted an epigenomically-conserved promoter in the *PRKCE* locus. In current GWAS catalogues, genetic associations in this locus include blood pressure regulation within the region we targeted ^45,46^. PRKCE is a member of protein kinase C family of proteins and has important roles in cardiomyocytes ^47^, with the epigenomically-conserved promoter associating with the longest isoform in this locus. Although variable across differentiation batches, we observed the expected reductions in *PRKCE* expression upon homozygous deletion of the target promoter in iPSC-cardiomyocytes, especially at day 15 post-differentiation (Figure 3C).

These results across epigenomically-conserved regulatory elements in three cardiovascular loci support their significant contributions to target gene expression, and constitute a cellular resource in which to explore their wider regulatory and cellular impact.

### Epigenomically-conserved promoters and enhancers impact global gene expression during cardiomyocyte differentiation

To further characterise gene expression contributions of epigenomically-conserved enhancers and promoters across the three targeted loci, we profiled genome-wide mRNA levels for wild- type and homozygous deletion iPSC clones at two time-points during cardiomyocyte differentiation.

We first asked whether gene expression changes upon genomic deletion of target sequences in the *KCNH2* locus may differ between epigenomically-conserved and control, human-only enhancers (Figure 4A, Figure S4 and Table S4). Comparison of differentially-expressed genes (DEGs) between homozygous deletions and wild-type controls for each enhancer revealed a much more pronounced effect on downstream gene expression for the epigenomically- conserved enhancer at 15 days of cardiomyocyte differentiation (Figure 4A); a difference that was similarly observed at the later time-point (Figure S4). These findings further support a strong contribution to downstream gene expression for the epigenomically-conserved *KCNH2* enhancer in differentiated cardiomyocytes, compared to negligible effects for the control, human-only enhancer.

**Figure 4:**
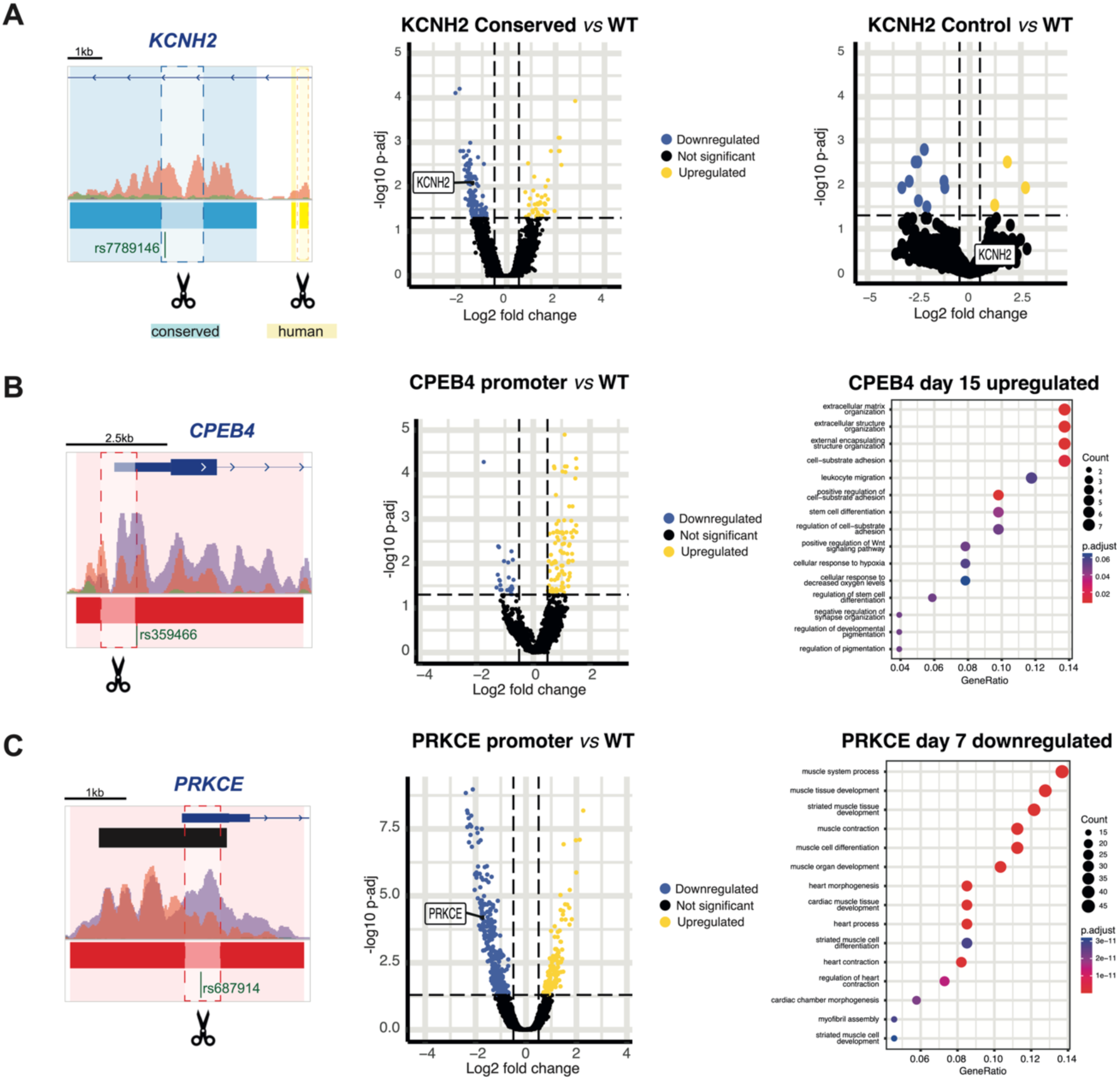
Gene expression changes associated to genomic deletion of candidate enhancer and promoter loci during cardiomyocyte differentiation **A.** Widespread gene expression changes upon deletion of an epigenomically-conserved enhancer in the *KCNH2* locus (blue; Conserved vs WT, middle), as compared to a control nearby enhancer (enhancer 2, yellow; right). The left diagram summarises epigenomic enrichments around the targeted enhancers and cardiovascular common genetic variants (details as in Figure 3). Volcano plots represent gene expression change (log2 fold changes; x-axis) and statistical significance (-log10(p-value)) for differentially expressed genes in each comparison (downregulated in dark blue and upregulated in yellow; Methods) at day 15 of cardiomyocyte differentiation. **B.** A promoter deletion in the *CPEB4* locus (left diagram) largely results in gene upregulation (volcano plot, middle) at day 15 of cardiomyocyte differentiation. Gene ontology enrichments for upregulated genes are represented as a circle plot (count (size) and adjusted p-value (shade); Methods). Details of diagram and volcano plot as in A. **C.** Gene expression changes across cardiomyocyte processes associate with deletion of a *PRKCE* candidate promoter at day 7 post-differentiation. Differentially expressed genes upon promoter deletion in the *PRKCE* locus (centre). Gene ontology (GO) enrichments for downregulated genes include heart biological processes (right). Details of diagram and volcano plot as in A, and GO enrichments as in B. See also **Figure S4**, **Table S4** and **Table S5**.

We next assessed downstream gene expression changes upon genomic deletion of epigenomically-conserved promoters in the *CPEB4* and *PRKCE* loci. Homozygous deletion of *CPEB4* promoter led to widespread gene expression changes at days 7 and 15 of differentiation, with a majority of DEGs being upregulated in the promoter knock-out samples at day 15 (Figure 4C,D and Figure S4). In contrast, genomic deletion of *PRKCE* promoter was associated with similar numbers of upregulated and downregulated genes, although gene expression changes were more pronounced in day 7 samples (Figure 4E,F and Figure S4).

These results indicate an important role for the two epigenomically-conserved promoters in gene expression regulation during cardiomyocyte differentiation. *CPEB4* promoter deletion typically led to gene upregulation, an observation that may relate to the roles of CPEB RNA binding proteins in promoting mRNA stability ^48^. Genomic deletion of *PRKCE* promoter associated both with upregulation and downregulation of genes, as expected for gene expression effects downstream to protein kinase signaling ^47^.

To further characterize the gene expression outcomes of promoter and enhancer deletions, we investigated their association of differentially expressed genes with gene ontology terms (Figure 4B-C, Figure S4, Table S5). As expected, and consistent with the known roles of KCNH2 in cardiac repolarization, genes downregulated upon deletion of the epigenomically- conserved enhancer in this locus were associated with ontology terms for heart biological processes at both time points (Figure S4D), including cardiac muscle contraction. In contrast, genes downregulated upon deletion of the epigenomically-conserved promoter in the *CPEB4* locus were enriched for stress pathway gene ontologies, including apoptosis and DNA damage (Figure 4B and Figure S4E); consistent with the roles of CPEB4 as a survival protein ^39^. However, most DEGs upon *CPEB4* promoter deletion were upregulated, and associated with ontology terms across cellular responses such as extracellular matrix organization and cell adhesion. Lastly, genes downregulated in cardiomyocytes derived from iPSCs with a *PRKCE* promoter deletion were enriched for gene ontologies of cardiac development, differentiation and contraction processes. This observation is in line with the functional role of PRKCE signalling in cardiomyocytes, particularly as part of sarcomeric contraction ^47^. However, upregulated genes upon KCNH2 enhancer or *PRKCE* promoter deletions tended to associate with general biological processes rather than those specific to cardiomyocytes (Figure S4D,F), suggesting they may correspond to gene expression changes secondary to KCNH2 and PRKCE downregulation.

In sum, gene expression changes upon deletion of three epigenomically-conserved enhancer and promoter candidates in differentiated cardiomyocytes support their significant contributions to gene expression. Moreover, epigenomically-conserved promoters in *CPEB4* and *PRKCE* loci associate with distinct ontology terms, consistent with biological processes relevant to cardiomyocytes and their responses to stress.

### The impact of epigenomically-conserved promoters of CPEB4 and PRKCE expression on cardiomyocyte phenotypes

The association of epigenomic conservation in the heart with trait-associated genetic variants and downstream gene regulation in cardiomyocytes (Figure 2) suggests this subset of regulatory elements may influence cellular phenotypes. Indeed, the epigenomically-conserved enhancer we targeted in the *KCNH2* locus (Figure 4A) has been previously shown to impact action potential duration in human iPSC-cardiomyocytes ^31^, consistent with genetic associations in this locus for QT interval duration and atrial fibrillation ^37,38,49^. To further test this hypothesis experimentally (Figures 3 and 4), we focused on candidate epigenomically- conserved promoters in the *CPEB4* and *PRKCE* loci, and selected two cellular phenotypes in heart muscle cells based on current genetic associations and literature evidence across these loci.

Genetic associations for *CPEB4* include myocardial mass and its associated electrocardiographic traits, such as the amplitude or duration of the QRS complex ^7^. Moreover, a recent cellular screening for RNA-binding proteins ^41^ suggested CPEB4 as a regulator of cardiomyocyte hypertrophy in neonatal rat cells. Therefore, we first interrogated the effect of *CPEB4* and *PRKCE* deletions on cellular hypertrophy in human iPSC-cardiomyocytes, by adapting a previously developed method ^24^ to quantify increased cellular size in response to the hypertrophic peptide endothelin-1 (ET-1) (Figure S5A). In this system, 72-hour treatment of wild-type cultures with ET-1 led to a significant increase in cardiomyocyte size of ∼25%, in line with previous observations in stem-cell derived human cardiomyocytes ^24,50,51^. In contrast, genomic deletion of the *CPEB4* candidate promoter led to negligible changes in cardiomyocyte size upon ET-1 treatment (Figure 5A and Figure S5B), albeit cell size in basal untreated cultures was consistently increased (as observed previously ^41^). These results suggest the epigenomically-conserved promoter in the *CPEB4* locus is necessary for a full hypertrophic response in human cardiomyocytes, with its deletion leading to hypertrophy in basal conditions and in the absence of ET-1 stimulation. In the same experimental system, a similar promoter deletion in the *PRKCE* locus did not associate to consistent differences in cellular size compared to wild-type cultures, either in basal conditions or following ET-1 treatment (Figure 5A and Figure S5B). To corroborate this observation, we additionally assessed expression of the hypertrophy-responsive genes *MYH7*, *ACTA1* and *NPPB* upon ET-1 treatment for wild-type control cardiomyocytes and those derived from *CPEB4* and *PRKCE* promoter deletion clones (Figure 5B and Figure S5B). Interestingly, the promoter *CPEB4* deletion abrogated induction of hypertrophy-responsive genes by ET-1 (Figure 5A and 5B), suggesting basal induction of hypertrophy mediated by reduced *CPEB4* expression associates with a poor cellular and gene expression response to ET-1. In contrast, for the *PRKCE* promoter deletion we observed a similar ET-1 response to wild-type cultures across hypertrophy-responsive genes, consistent with a modest impact of this promoter deletion on cardiomyocyte hypertrophy *in vitro*.

**Figure 5:**
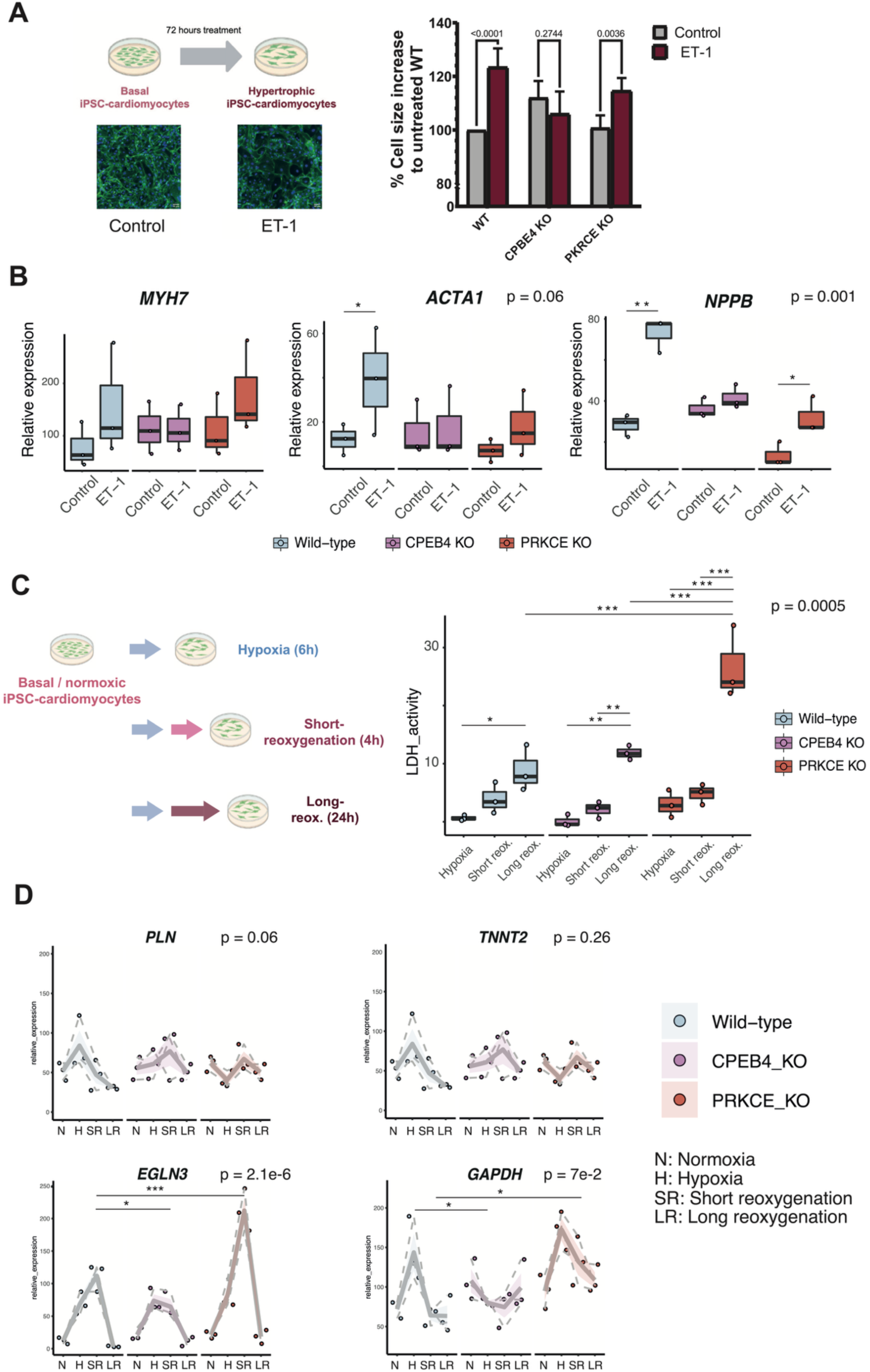
Examination of the impact of candidate promoter deletions on cardiomyocyte hypertrophy and sensitivity to hypoxia-reoxygenation **A.** Increase of cardiomyocyte size in response to endothelin-1 is dependent on an epigenomically-conserved promoter in the *CPEB4* locus, but not on a similar deletion in the *PRKCE* locus. Changes in cardiomyocyte size were measured in day 18 cultures in response to treatment with endothelin-1, and quantified with high-content microscopy (left diagram, 50 micron scale bar for example images, Methods). Percentage changes in size are compared between control and ET-1 treated cultures (barplot, right) across the following genotypes: wild- type, *CPEB4* promoter knock-out (CPEB4 KO) and *PRKCE* promoter knock-out (PRKCE KO). Data corresponds to imaging in three independent batches of iPSC-cardiomyocyte differentiation. P-values: two-way ANOVA with Tukey post-hoc correction (Methods). **B.** Gene expression levels of hypertrophy markers *MYH7*, *ACTA1* and *NPPB* were quantified wild-type (light blue), *CPEB4* promoter KO (violet) and *PRKCE* promoter KO (red). P-values above each plot correspond to two-way repeated measures ANOVA for interactions between treatment and genotype. Number of independent measurements as in A. Pairwise comparisons: Tukey post-hoc correction: ** < 0.01, * < 0.05 **C.** Cytotoxicity induced by hypoxia-reoxygenation treatment in day 22 cardiomyocytes. Lactate dehydrogenase (LDH) activity released to culture media was measured to quantify cytotoxicity in iPSC cardiomyocyte cultures after 6 hours of hypoxia (Hypoxia), or the same exposure followed by either 4 hours (Short reox.) or 24 hours of reoxygenation (Long reox.); and is represented relative to maximum activity (Methods) across the three genotypes (details as in B). Number of independent measurements as in A. Two-way ANOVA p-value=0.0005 for interaction between treatment and genotype. Pairwise comparisons: Tukey post-hoc correction: *** < 0.001, ** < 0.01 **D.** Gene expression of cardiac (*PLN* and *TNNT2*) and hypoxia-responsive markers (*EGLN3* and *GAPDH*) were quantified in hypoxia-reoxygenation samples from day XX cardiomyocytes across the three genotypes (details as in C). Data is represented as gene expression trendlines from triplicate samples across genotypes and treatments. Number of independent measurements as in A. P-values: Two-way repeated measures ANOVA for interaction between treatment and genotype; Tukey post-hoc correction for pairwise p-values *** < 0.001, * < 0.1 See also **Figure S5**.

To further investigate the impact of the either promoter deletion on cellular phenotypes, we selected a second cellular response in cardiomyocytes. Cardiovascular genetic associations in the PRKCE locus are varied and include multifactorial traits such as diastolic blood pressure ^45^, and previous reports identified *PRKCE* as a regulator of cardiomyocyte necrosis ^52^. Based on these observations, we assessed the impact of *PRKCE* and *CPEB4* promoter deletions on the response of iPSC-cardiomyocytes to hypoxia, and either short or long periods of reoxygenation (Figure 5C). This is a standard experimental system to investigate cytotoxicity in cardiomyocytes ^25^, in which we measured cardiac, hypoxia-responsive ^53^ and reoxygenation-responsive ^54^ genes (Figure S5C, Methods). This experiment showed deletion of *PRKCE* promoter was associated with significantly increased cytotoxicity in cardiomyocytes compared to wild-type controls, whereas the *CPEB4* promoter deletion had a non-significant effect (Figure 5C). For *PRKCE,* these results are consistent with previous biochemical evidence of increased cardiomyocyte cytotoxicity upon treatment with a selective PRKCE inhibitor ^55^. To investigate potential mechanisms underlying the observed changes in cytotoxicity, we assessed gene expression across cardiac-specific genes *PLN*, *TNNT2* and *MYH7*, as well as hypoxia-responsive genes *EGLN3*, *GAPDH* and *GYS1* – which are known targets of hypoxia-inducible factor ^53,56–58^. We did not observe significant changes in the expression of cardiac-specific genes across wild-type, *CPEB4*- or *PRKCE*-KO cardiomyocytes (Figure 5D and Figure S5C). In contrast, induction dynamics for hypoxia- responsive genes were altered upon deletion of *CPEB4* or *PRKCE* promoters. The first led to a delay in *EGNL3* returning to basal levels in the reoxygenation phase, and increased overall levels of *GAPDH* and *GYS1*; whereas deletion of the *PRKCE* promoter resulted in increased gene expression of hypoxia-responsive gene, especially during short reoxygenation. Although limited by the small set of genes we evaluated, these observations suggest reductions in *CPEB4* or *PRKCE* expression alter the dynamics of the hypoxia- reoxygenation response in cardiomyocytes, potentially through direct or indirect effects on mRNA stability. In contrast, we did not observe consistent changes for the expression dynamics of reoxygenation-responsive genes (Figure S5D), for which we used a panel of NRF2 target genes ^54^.

In sum, these results substantiate our genome-wide findings on how epigenomic conservation in the heart enriches for trait-associated genetic variation and downstream gene regulation in cardiomyocytes; and suggest this subset of regulatory regions is more likely to impact downstream cellular phenotypes.

## DISCUSSION

Vast improvements in statistical power for human population studies are increasingly uncovering a complex ensemble of trait- and disease-associated genetic variation occurring in non-coding regions of the genome ^2,5,6^. As such, epigenomic annotation of genetic association loci is commonly used to support fine-mapping and functional evaluation efforts ^59,60^. From an evolutionary perspective, comparative epigenomic studies across species and developmental contexts have documented a continuum of regulatory regions from highly-conserved to lineage-specific ^14,15,61^. The first has been associated with regulatory pleiotropy and core tissue functions ^16,33,34^, whereas lineage-specific changes in regulatory activity are thought to contribute – at least in some cases – to tissue-level phenotypic adaptation ^29,62^. However, the extent to which trait-associated genetic variation may preferentially occur in highly-conserved or lineage-specific regulatory regions remains largely unexplored, with the exception of limited comparisons in primate tissues and *in vitro* models ^17,20^.

Outside a subset of well-characterised examples (reviewed in ^8^), the regulatory and phenotypic impact of most non-coding genetic variation remains unexplored. Firstly, most lead variants are in strong linkage disequilibrium with many other positions, and it remains challenging to prioritise variants for experimental perturbation experiments ^63^. Secondly, the effect sizes for most common trait-associated genetic variants are small, and experimentally interrogated variants often have a limited effect on transcriptomic or cellular readouts ^64^. An alternative approach relies on targeting regulatory elements harboring trait- and disease-associated variants for functional evaluation studies ^65,66^, which often have measurable effects in experimental *in vitro* systems ^67^. To investigate the interplay between epigenomic regulatory conservation and trait-associated genetic variation, here we combined comparative epigenomics of heart tissue samples across mammals with experimental *in vitro* perturbations in human iPSC-derived cardiomyocytes. This strategy allowed us to assess the distribution of cardiovascular common genetic variation across highly-conserved and lineage-specific promoters and enhancers in the heart, select candidate trait-associated regulatory elements for perturbation experiments, and assess transcriptional and cellular effects downstream of targeted candidate elements across three cardiovascular loci. Our work adds to ongoing efforts to select and prioritise disease-associated regulatory elements for interrogation of their regulatory and phenotypic impact ^8,63^.

Our results in heart tissue further support an increased functional relevance for highly-conserved regulatory elements and their association with cardiovascular trait-associated genetic variation. These observations extend previous results in primate tissues ^17^, for which comparison of human and marmoset epigenomes similarly found disease-associated variants enriched within conserved regulatory activity. Moreover, we found epigenomically- conserved promoters and enhancers in the heart associate with signatures of gene regulation across multiple heart cell types. The first agrees with previous work across mammalian tissues ^16,33^ associating regulatory pleiotropy with mammalian epigenomic conservation, which we observe in heart tissue across major cardiac cell types. Our results also suggest pleiotropic genetic variation may associate with highly-conserved regulatory elements, as proposed previously in other tissues and contexts ^34,68^. For cardiomyocytes, our results for the two candidate enhancers in the *KCNH2* locus are consistent with earlier observations across mammalian tissues indicative of stronger gene expression contributions for epigenomically-conserved regulatory activity ^16,18,19^.

Comparing cardiomyocyte responses to hypertrophic and oxidative stress for candidate promoter deletions in the *CPEB4* and *PRKCE loci*, our results inform the cellular outcomes upon perturbation of epigenomically-conserved promoter activity. We found cardiomyocyte hypertrophy is significantly altered by deletion of a highly-conserved promoter in *CPEB4*, in agreement with previous reports in neonatal rat cardiomyocytes ^41^. As an RNA-binding protein, CPEB4 may regulate mRNA stability of transcripts during cardiomyocyte hypertrophy, as suggested by our observation of reduced induction of hypertrophy markers by ET-1. In contrast, a similar promoter deletion in the *PRKCE* locus did not significantly impair the response to hypertrophic stimuli, and had only a modest effect on the induction of hypertrophy marker genes. However, in response to hypoxia/reoxygenation, we observed a significant increase in cytotoxicity upon deletion of this promoter region. This observation aligns with previous biochemical and cellular evidence implicating PRKCE loss-of-function in cardiomyocyte cytotoxicity and necrosis ^52,55^. By measuring gene expression of selected cardiac, hypoxic and reoxygenation markers, our results potentially implicate the candidate *CPEB4* and *PRKCE* promoter deletions in regulating the dynamics of hypoxia-reoxygenation responses, as suggested by increased expression levels of known HIF-target genes upon promoter perturbations. Further experimental work will be required to elucidate the molecular mechanisms underlying our observations. However, these may relate to the proposed role of CPEB family members in regulation of mRNA stability ^39^ or HIF translation ^40^, whereas the effects we observed upon *PRKCE* promoter deletion may be a consequence of its effects in cardiomyocyte viability. On the whole, our results across candidate loci support the relevance of epigenomically-conserved regulatory activity in the heart in regulation of gene expression and cardiomyocyte phenotypes.

There are a number of limitations to our approach. First, by profiling and experimentally targeting epigenomically-conserved promoter and enhancer regions, our approach cannot directly assess the contribution of common genetic variants within candidate elements. As expected, and compared to genome editing of individual genomic variants, deletion of epigenomically-conserved promoters or enhancers in our study showed effects on gene expression and cellular responses consistent with moderate effect sizes ^19^. Nevertheless, further experimental work would be required to functionally interpret the impact of individual genetic variants on the activity of our candidate regulatory elements.

Second, our experimental perturbation approach relied on deletion of individual epigenomically-conserved enhancers and promoters defined from canonical histone marks ^69,70^ which have been previously proposed to strongly contribute to downstream gene expression ^19,33,34^. However, the substantial complexity of epigenomic signals in most trait-associated loci will increasingly demand approaches amenable to combinatorial perturbations ^66,71^, which can include interrogation of non-canonical epigenomic signals as an additional source of regulatory potential ^72,73^.

Third, our approach focused on comparative genomics in heart tissue and perturbation experiments in human cardiomyocytes. This choice is informed by the relative under- representation of the heart in previous comparative approaches ^16,17^, and the strong heart association of genetic association signals across a range of human traits and diseases ^2^. Nevertheless, our results also suggest epigenomic enhancer conservation is associated with pleiotropic regulatory properties, such as chromatin accessibility across several cardiac cell types ^31^. The candidate epigenomically-conserved promoters and enhancers we defined here are thus likely to impact gene expression and cellular functions in additional heart cell types, and potentially in other somatic tissues.

In summary, this study presents a comparative epigenomics angle with which to investigate the regulatory contribution of trait-associated genetic variation in the heart. By integrating epigenomic, transcriptomic and cell biology analyses, we show the utility of this approach in delineating trait-associated promoters and enhancers with conserved regulatory activity, and experimentally connect them with cardiomyocyte-specific gene expression and cellular responses.

## MATERIALS AND METHODS

### Chromatin immunoprecipitation and high-throughput sequencing *– related to Figure 1*

We performed chromatin immunoprecipitation experiments followed by high throughput sequencing (ChIP-seq) using heart tissue samples isolated from rhesus macaque, marmoset, mouse, rat, rabbit, cat, dog, horse, cow and pig. The origin, number of replicates, sex, and age for each species’ samples are detailed in Table S1 and ArrayExpress accession E-MTAB- 11559. For the human heart, we used previously-reported datasets ^2^.

All animal experiments were conducted in accordance with the UK Animals (Scientific Procedures) Act 1986 Amendment Regulations 2012 and performed under the terms and conditions of UK Home Office licenses P51E67724 (D.V.) and PP8702170 (D.V.). organ in small pieces around 1cm^3^. Blood clots within the heart ventricles were removed. Cross-linking of the diced tissue was performed in 1% formaldehyde solution for 20 min, addition of 250 mM glycine and incubation for a further 10 min to neutralize the formaldehyde. After homogenization of cross-linked tissues in a dounce tissue grinder, samples were washed twice with PBS and lysed according to published protocols ^75^ to solubilize DNA-protein complexes. Chromatin was fragmented to 300 bp average size by sonication on a Misonix sonicator 3000 with a 418 tip (1/16 inch diameter). Chromatin from 50-200 mg of dounced tissue was used for each ChIP experiment using antibodies against H3K4me3 (millipore 05- 1339), H3K27ac (abcam ab4729) and H3K4me1 (abcam ab8895). Illumina sequencing libraries were prepared from ChIP-enriched DNA using ThruPLEX DNA-seq library preparation kit (Takara Bio) with up to 10ng of input DNA and 8-15 PCR cycles. After PCR, libraries were pooled in equimolar concentrations and sequenced on Illumina HiSeq 4000 or NovaSeq instruments.

For the purpose of epigenomic annotation of candidate promoters and enhancers *towards in vitro* experiments, ChIP-seq was also carried out from iPSC-cardiomyocytes at day 20 post-differentiation (following metabolic selection ^76^).

### Computational analysis of ChIP-seq data – related to Figure 1

<u>Basic alignment and peak calling</u>: Aligned bam files were obtained with bwa 0.7.17 and the Ensembl v99 assemblies indicated in Table S1. We used macs2 ^77^ to call peaks for each ChiP- Seq replicate, using default parameters and “--keep-dup all” to retain duplicate reads. Before peak-calling, multi-mapping reads were removed and read-depth adjusted to 20 million uniquely mapped reads (or all available reads for low-depth libraries).

<u>Definition of regulatory regions from ChIP-seq peaks</u>: We first constructed sets of reproducible peaks for each combination of histone mark, tissue and species by merging peaks identified across a minimum of two biological replicates (with minimum 50% length overlap). In a second step, we used these sets of reproducible peaks to identify promoters, enhancers and primed enhancers independently for each species and tissue, according to the following criteria: all H3K4me3 peaks overlapping an H3K27ac peak (minimum 50% bases overlap) were predicted to be promoters, H3K27ac peaks not overlapping promoters were predicted as enhancers, and H3K4me1 peaks not overlapping any H3K4me3 or H3K27ac peaks were as predicted primed enhancers.

### Cross-mapping of regulatory regions across 11 mammalian species

<u>Definition of orthologous regulatory regions</u>: we used the LiftOver local software ^78^ to map genomic coordinates of the regulatory elements identified in each species to the human genome. LastZ pairwise whole-genome alignments with human were downloaded from Ensembl Compara v99 and converted from ensembl maf format to the UCSC chain format using UCSC tools (including mafToPsl, pslToChain and chainSwap, v357 for all software). We validated the correct implementation of the coordinates conversion step using LiftOver with the generated chain alignment files by comparing the resulting coordinates with those obtained from queries directly performed through the Ensembl API^79^.

We defined a set of “high-confidence orthologous regions” as 11-way orthologous regions for which we required robust LiftOver mapping of regulatory elements across the study genomes. This was achieved in two steps, involving (i) the definition of the set of regulatory regions, expressed in human genome coordinates, that can be aligned from each of the other species to human; and (ii) a filtering step to retain only (sub)regions with a strict reciprocal LiftOver mapping in each genome (e.g. a marmoset region mapping to human, and the human coordinates of this region mapping back to each of the other 10 genomes). In this filtering step, the final criteria for regions to be defined as high-confidence 11-way orthologs were: a reciprocal LiftOver mapping from human to the ther genomes, with a minimum of 10% coverage (-minMatch 0.10) and similar lengths of the resulting regions across the study genomes (the difference in length between the human region and regions in any other species must be less than 15% of the largest region).

<u>Homogenization of regulatory region type</u>: for cases where orthologous regions were defined as different types of regulatory elements across species, we used a majority rule system to homogenize regulatory element types. For instance, a region defined as a promoter in human, marmoset and cow but as an enhancer in horse was re-defined as a promoter across all species. In case of ties, regions are arbitrarily assigned to the “highest-level” regulatory type (i.e. promoters have priority over enhancers and primed enhancers, and enhancers have priority over primed enhancers).

### Computational analyses of GWAS variants and signatures of gene regulation – related to Figure 2

Genetic lead variants associated to cardiovascular traits at genome-wide significance (p-value < 10^-8^) were obtained from the NHGRI-EBI GWAS catalog ^6^ (version e113, 2025), for all heart-related traits (search term “heart” across associated traits in the catalog), and four major cardiovascular diseases and traits (search terms “atrial fibrillation, “QT interval”, “coronary artery disease” and “cardiomyopathy”). Each set of lead variants was annotated with proxy variants in strong LD (R^2^ > 0.8) using SNIPA ^80^. The enrichment of GWAS SNPs in heart promoters and enhancers across epigenomically-conserved, primate-specific and human only categories calculated with a hypergeometric test.

Signatures of gene regulation across heart cell types and cardiomyocytes were obtained from previously published datasets: RPKM chromatin accessibility levels across cardiac cell types ^31^, and significant promoter-enhancer interactions in human iPSCs and iPSC-derived cardiomyocytes were obtained from ^32^.

### Selection of candidate regions for genetic deletion experiments – related to Figures 2 and 3

Candidate promoters and enhancers for genetic deletion experiments were selected according to the following selection criteria (see also Figure S3A): epigenomic conservation across study species (and thus classified as “epigenomically-conserved”), proximity to cardiovascular GWAS SNPs (within 1 Kb window), evidence of conserved regulatory activity in iPSC-cardiomyocytes and human left ventricle ^2^, overlap with significant promoter-enhancer interactions in iPSC-cardiomyocytes ^32^, and containing one or more single-cell ATAC-seq peaks corresponding to accessible chromatin regions in human cardiomyocytes ^31^. For CRISPR/Cas9 genomic deletion, we excluded candidate regions overlapping coding exons, or those with poor guide RNA design metrics.

### Genomic deletion of candidate regulatory regions in human iPSCs – related to Figures 3, 4 and 5

CRISPR guide RNA (gRNA) were selected using CHOPCHOP ^81^. A gRNA was selected to target each end of genomic regions of interest with best combination of positioning, efficiency score and minimal predicted off-targets ^82^. gRNA-Cas9 plasmids were created through modification of pSpCas9(BB)-2A-GFP plasmid (PX458, Addgene #48138; ^23^). The GFP reporter was replaced through subcloning either by mCherry (Addgene# 154193) or BFP sequences (Addgene#60955) to create hU6-gRNA-Cbh-Cas9-T2A-mCherry and hU6-gRNA-Cbh-Cas9-T2A-BFP, respectively. These plasmids were digested using BbsI before ligation with phosphorylated double-stranded oligonucleotides ^23^. For each target genomic region, one guide was cloned into the mCherry plasmid, and the second into the BFP plasmid; guide RNA sequences are detailed in Table S6.

GenC WTC/GM25256 iPSCs ^83^ were cultured on recombinant human vitronectin (VTN-N; Gibco) with mTeSR Plus media (STEMCELL Technologies). Cells were regularly passaged at 70-80% confluency using Versene (Gibco). During, and 24hr post passaging/seeding, media was supplemented with 10µM Y-27632 (Abcam).

iPSC (2.5x104 cells per cm2) were reverse transfected with 150ng plasmid/cm2 of each gRNA-Cas9-BFP/mCherry plasmid using Lipofectamine Stem Transfection Reagent (Invitrogen) at 3:1 ratio. 48hr post-transfection, cells were sorted using FACSAria Cell Sorter (BD Biosciences). BFP and mCherry double-positive cells were seeded at single cell density in mTeSR Plus supplemented with CloneR2 (STEMCELL Technologies). Single, clonal colonies were propagated for further experiments. Clone genotype was verified using PCR and Sanger Sequencing (Genewiz), using sequencing primers in Table S6.

### Cardiomyocyte differentiation from human iPSCs – related to Figures 3, 4 and 5

iPSC were differentiated into cardiomyocytes using GiWi protocol ^21^. Two days prior to induction of differentiation, iPSC were seeded at 7.5x10^4^ cells per cm^2^ on hESC-Qualified Geltrex (Gibco) coated plates. On day 0, with cellular confluency of ∼90%, media was replaced with cardiac differentiation media (RPMI 1640 (Corning) and B-27 minus insulin (Gibco)), containing 6µM CHIR99021 (AbMole). On day 2, media was changed to cardiac differentiation media containing 2µM Wnt-C59 (AdooQ Bioscience). From day 4, cardiac differentiation media was changed every 48hr. From day 7 onwards, B-27 (Gibco) was used in media. Between days 12 and 15, cells were cultured in lactate selection media ^76,84^ before return to cardiac differentiation media until sample collection.

### RT-qPCR – related to Figures 3, 4 and 5

RNA was extracted using Monarch Total RNA Miniprep kit (New England BioLabs). 1µg RNA was reverse transcribed using High-Capacity cDNA Reverse Transcription Kit (Applied Biosystems). RT-qPCR was carried out with 5ng cDNA per 10µL reaction using with KAPA SYBR FAST qPCR Universal (KAPA Biosystems) on Roche LightCycler® 480 (Roche), using primer sequences detailed in Table S7. *TBP* was used as an internal reference for gene expression normalisation.

### RNA-sequencing experiments and analyses – related to Figure 4

RNA integrity was assessed using RNA 6000 Nano Kit for Bioanalyzer 2100 (Agilent). RNA-seq libraries were prepared using New England BioLabs NEBNext workflow. Each step was carried out in accordance with manufacturer’s guidelines; 1000ng of total RNA was rRNA depleted using NEBNext rRNA Depletion Kit v2 before preparation of libraries using NEBNext Ultra II Directional RNA Library Prep Kit for Illumina and NEBNext Multiplex Oligos for Illumina (Dual Index Primers Set 1). DNA bead purification steps were carried out using AMPure XP beads (Beckman Coulter).

Final library concentrations were calculated via qPCR using KAPA Library Quantification Kit (Roche). Libraries were pooled at equimolar final concentration before PE150 Illumina sequencing in NovaSeq instruments (Novogene, UK).

### Hypertrophy induction and high-content microscopy quantification of cardiomyocyte size – related to Figure 5

0.5x10^6^ iPSC-CMs per well were replated at day 10 in a 6-well plate, and media refreshed every 48h. After allowing cells to recover, cells were treated with 10nM endothelin-1 (Tocris) for 72h. Then cells were fixed with 4% PFA for 10 minutes at room temperature, after 2x10min washings with PBS, cells were stained with 10μg/ml WGA-FITC (Generon) for 10 minutes at room temperature, cells were washed again 2x10min with PBS and stained with 1mg/mL Hoechst (Fisher scientific) for 1 minute. Lastly, cells were extensively washed (4x10min) and 24 images per well were acquired using the INCA6000 (GE) automated confocal microscope. Images were first subjected to machine-learning cell segmentation by using *ilastik* ^85^ and subsequently cell area was measured with CellProfiler 4.2.1.

### Exposure to hypoxia/reoxygenation and cellular cytotoxicity analysis – related to Figure 5

1x10^6^ iPSC-CMs were replated at day 10, with lactate selection ^84^ started at day 14 over 4 days (refreshing media every 48h). Media was changed to cardiac differentiation media with B27 supplement (Gibco) and refreshed every 48h. Hypoxia/reoxygenation treatments were started at the indicated time points in a H35 HEPA Hypoxia workstation (Don Whitley scientific), as follows: hypoxia (6 hours incubation at 1% O2, 5%CO2), short reoxygenation (6 hours incubation at 1%O2, 5%CO2 followed by 4 hours of reoxygenation at 21%O2, 5%CO2), or long reoxygenation (6 hours incubation at 1%O2, 5%CO2 followed by 24h reoxygenation at 21% O2, 5%CO2).

For each replicate experiment, cell cultures for each treatment were staggered in advance so all treatments finished at the same differentiation time point (day 22), thus allowing simultaneous sample collection and measurements.

Lactate dehydrogenase (LDH) activity was measured with CyQUANT LDH Cytotoxicity Assay Kit (Invitrogen) using 50uL of sample medium and following manufacturer instructions. Absorbance at 490nm and 680nm (background) was measured in a FLUOstar Omega plate reader (BMG Labtech). Percentage of LDH activity was calculated with the following formula:

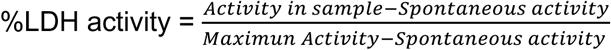

Activity in sample corresponds to absorbance in culture media for each sample, spontaneous activity to absorbance in untreated controls, and maximum activity to absorbance in cell lysates.

## AUTHOR CONTRIBUTIONS

SF, DFA, AR, DPH and DV designed and performed experiments. SF, EP, DFA and DV analysed data. AT provided essential reagents and technical advice. DV and DFA secured funding. DV coordinated the study and wrote the manuscript. All authors read and approved the manuscript.

## ACKNOWLEDGEMENTS

This work has been funded by the British Heart Foundation (FS/18/39/33684 to D.V.), Barts Charity (Seed grant G-002635 to D.V. and D.F.A) and Queen Mary University of London. DFA was supported by a personal Fellowship from Alfonso Martin Escudero Foundation, and AR by a PhD fellowship grant from Barts Charity (MGU0501 to D.V.). We thank Camille Berthelot (Institut Pasteur, Paris), Luis del Peso (Medicine School, UAM, Madrid) and members of the QMUL Centre of Epigenetics for critical advice and suggestions, and Drs. Luke Gammon and Gary Warnes (Blizard Institute) for technical assistance.

## DATA AVAILABILITY

The ChIP-sequencing data reported here across mammalian species has been deposited to ArrayExpress with accession number E-MTAB-11559. RNA-sequencing and ChIP-sequencing datasets in human iPSC-cardiomyocytes have been deposited to GEO (GSE287934 and GSE288112, respectively).

## REFERENCES

1 Spitz, F. & Furlong, E. E. Transcription factors: from enhancer binding to developmental control. Nat Rev Genet 13, 613–626, doi:10.1038/nrg3207 (2012).

2 Roadmap Epigenomics, C., et al. Integrative analysis of 111 reference human epigenomes. Nature 518, 317–330, doi:10.1038/nature14248 (2015).

3 Claringbould, A. & Zaugg, J. B. Enhancers in disease: molecular basis and emerging treatment strategies. Trends Mol Med 27, 1060–1073, doi:10.1016/j.molmed.2021.07.012 (2021).

4 Smemo, S. et al. Regulatory variation in a TBX5 enhancer leads to isolated congenital heart disease. Hum Mol Genet 21, 3255–3263, doi:10.1093/hmg/dds165 (2012).

5 Maurano, M. T. et al. Systematic localization of common disease-associated variation in regulatory DNA. Science 337, 1190–1195, doi:10.1126/science.1222794 (2012).

6 Cerezo, M. et al. The NHGRI-EBI GWAS Catalog: standards for reusability, sustainability and diversity. Nucleic Acids Res 53, D998–D1005, doi:10.1093/nar/gkae1070 (2025).

7 van der Harst, P. et al. 52 Genetic Loci Influencing Myocardial Mass. J Am Coll Cardiol 68, 1435–1448, doi:10.1016/j.jacc.2016.07.729 (2016).

8 Villar, D., Frost, S., Deloukas, P. & Tinker, A. The contribution of non-coding regulatory elements to cardiovascular disease. Open Biol 10, 200088, doi:10.1098/rsob.200088 (2020).

9 Jonker, T., Barnett, P., Boink, G. J. J. & Christoffels, V. M. Role of Genetic Variation in Transcriptional Regulatory Elements in Heart Rhythm. Cells 13, doi:10.3390/cells13010004 (2023).

10 Bezzina, C. R. et al. Common variants at SCN5A-SCN10A and HEY2 are associated with Brugada syndrome, a rare disease with high risk of sudden cardiac death. Nat Genet 45, 1044–1049, doi:10.1038/ng.2712 (2013).

11 van den Boogaard, M., et al. Genetic variation in T-box binding element functionally affects SCN5A/SCN10A enhancer. J Clin Invest 122, 2519–2530, doi:10.1172/JCI62613 (2012).

12 Ye, J. et al. A Functional Variant Associated with Atrial Fibrillation Regulates PITX2c Expression through TFAP2a. Am J Hum Genet 99, 1281–1291, doi:10.1016/j.ajhg.2016.10.001 (2016).

13 Cotney, J. et al. The evolution of lineage-specific regulatory activities in the human embryonic limb. Cell 154, 185–196, doi:10.1016/j.cell.2013.05.056 (2013).

14 Villar, D. et al. Enhancer evolution across 20 mammalian species. Cell 160, 554–566, doi:10.1016/j.cell.2015.01.006 (2015).

15 Vierstra, J. et al. Mouse regulatory DNA landscapes reveal global principles of cis-regulatory evolution. Science 346, 1007–1012, doi:10.1126/science.1246426 (2014).

16 Roller, M. et al. LINE retrotransposons characterize mammalian tissue-specific and evolutionarily dynamic regulatory regions. Genome Biol 22, 62, doi:10.1186/s13059-021-02260-y (2021).

17 Castelijns, B. et al. Recently Evolved Enhancers Emerge with High Interindividual Variability and Less Frequently Associate with Disease. Cell Rep 31, 107799, doi:10.1016/j.celrep.2020.107799 (2020).

18 Trizzino, M. et al. Transposable elements are the primary source of novelty in primate gene regulation. Genome Res 27, 1623–1633, doi:10.1101/gr.218149.116 (2017).

19 Berthelot, C., Villar, D., Horvath, J. E., Odom, D. T. & Flicek, P. Complexity and conservation of regulatory landscapes underlie evolutionary resilience of mammalian gene expression. Nat Ecol Evol 2, 152–163, doi:10.1038/s41559-017-0377-2 (2018).

20 Destici, E. et al. Human-gained heart enhancers are associated with species-specific cardiac attributes. Nat Cardiovasc Res 1, 830–843, doi:10.1038/s44161-022-00124-7 (2022).

21 Lian, X. et al. Directed cardiomyocyte differentiation from human pluripotent stem cells by modulating Wnt/beta-catenin signaling under fully defined conditions. Nat Protoc 8, 162–175, doi:10.1038/nprot.2012.150 (2013).

22 Burridge, P. W. et al. Chemically defined generation of human cardiomyocytes. Nat Methods 11, 855–860, doi:10.1038/nmeth.2999 (2014).

23 Ran, F. A. et al. Genome engineering using the CRISPR-Cas9 system. Nat Protoc 8, 2281–2308, doi:10.1038/nprot.2013.143 (2013).

24 Ovchinnikova, E. et al. Modeling Human Cardiac Hypertrophy in Stem Cell-Derived Cardiomyocytes. Stem Cell Reports 10, 794–807, doi:10.1016/j.stemcr.2018.01.016 (2018).

25 Ward, M. C. & Gilad, Y. A generally conserved response to hypoxia in iPSC-derived cardiomyocytes from humans and chimpanzees. Elife 8, doi:10.7554/eLife.42374 (2019).

26 Harrison, P. W. et al. Ensembl 2024. Nucleic Acids Res 521, D891–D899, doi:10.1093/nar/gkad1049 (2024).

27 Desjardins, C. A. & Naya, F. J. Antagonistic regulation of cell-cycle and differentiation gene programs in neonatal cardiomyocytes by homologous MEF2 transcription factors. J Biol Chem 292, 10613–10629, doi:10.1074/jbc.M117.776153 (2017).

28 Kim, Y. et al. The MEF2D transcription factor mediates stress-dependent cardiac remodeling in mice. J Clin Invest 118, 124–132, doi:10.1172/JCI33255 (2008).

29 Parey, E. et al. Phylogenetic modeling of enhancer shifts in African mole-rats reveals regulatory changes associated with tissue-specific traits. bioRxiv, doi:10.1101/2023.01.10.523217 (2023).

30 Consortium, G. T. The GTEx Consortium atlas of genetic regulatory effects across human tissues. Science 369, 1318–1330, doi:10.1126/science.aaz1776 (2020).

31 Hocker, J. D. et al. Cardiac cell type-specific gene regulatory programs and disease risk association. Sci Adv 7, doi:10.1126/sciadv.abf1444 (2021).

32 Montefiori, L. E. et al. A promoter interaction map for cardiovascular disease genetics. Elife 7, doi:10.7554/eLife.35788 (2018).

33 Kliesmete, Z. et al. Evidence for compensatory evolution within pleiotropic regulatory elements. Genome Res 34, 1528–1539, doi:10.1101/gr.279001.124 (2024).

34 Fish, A., Chen, L. & Capra, J. A. Gene Regulatory Enhancers with Evolutionarily Conserved Activity Are More Pleiotropic than Those with Species-Specific Activity. Genome Biol Evol 9, 2615–2625, doi:10.1093/gbe/evx194 (2017).

35 Lyra-Leite, D. M. et al. A review of protocols for human iPSC culture, cardiac differentiation, subtype-specification, maturation, and direct reprogramming. STAR Protoc 3, 101560, doi:10.1016/j.xpro.2022.101560 (2022).

36 van den Boogaard, M., et al. Identification and Characterization of a Transcribed Distal Enhancer Involved in Cardiac Kcnh2 Regulation. Cell Rep 28, 2704–2714 e2705, doi:10.1016/j.celrep.2019.08.007 (2019).

37 Nielsen, J. B. et al. Biobank-driven genomic discovery yields new insight into atrial fibrillation biology. Nat Genet 50, 1234–1239, doi:10.1038/s41588-018-0171-3 (2018).

38 Roselli, C. et al. Multi-ethnic genome-wide association study for atrial fibrillation. Nat Genet 50, 1225–1233, doi:10.1038/s41588-018-0133-9 (2018).

39 Kan, M. C. et al. CPEB4 is a cell survival protein retained in the nucleus upon ischemia or endoplasmic reticulum calcium depletion. Mol Cell Biol 30, 5658–5671, doi:10.1128/MCB.00716-10 (2010).

40 Masuda, K., Abdelmohsen, K. & Gorospe, M. RNA-binding proteins implicated in the hypoxic response. J Cell Mol Med 13, 2759–2769, doi:10.1111/j.1582-4934.2009.00842.x (2009).

41 Riechert, E. et al. Identification of dynamic RNA-binding proteins uncovers a Cpeb4- controlled regulatory cascade during pathological cell growth of cardiomyocytes. Cell Rep 35, 109100, doi:10.1016/j.celrep.2021.109100 (2021).

42 Hoffmann, T. J. et al. Genome-wide association analyses using electronic health records identify new loci influencing blood pressure variation. Nat Genet 49, 54–64, doi:10.1038/ng.3715 (2017).

43 Sakaue, S. et al. A cross-population atlas of genetic associations for 220 human phenotypes. Nat Genet 53, 1415–1424, doi:10.1038/s41588-021-00931-x (2021).

44 Verma, A. et al. Diversity and scale: Genetic architecture of 2068 traits in the VA Million Veteran Program. Science 385, eadj1182, doi:10.1126/science.adj1182 (2024).

45 Warren, H. R. et al. Genome-wide association analysis identifies novel blood pressure loci and offers biological insights into cardiovascular risk. Nat Genet 49, 403–415, doi:10.1038/ng.3768 (2017).

46 Giri, A. et al. Trans-ethnic association study of blood pressure determinants in over 750,000 individuals. Nat Genet 51, 51–62, doi:10.1038/s41588-018-0303-9 (2019).

47 Scruggs, S. B., Wang, D. & Ping, P. PRKCE gene encoding protein kinase C-epsilon- Dual roles at sarcomeres and mitochondria in cardiomyocytes. Gene 590, 90–96, doi:10.1016/j.gene.2016.06.016 (2016).

48 Richter, J. D. CPEB: a life in translation. Trends Biochem Sci 32, 279–285, doi:10.1016/j.tibs.2007.04.004 (2007).

49 Newton-Cheh, C. et al. Common genetic variation in KCNH2 is associated with QT interval duration: the Framingham Heart Study. Circulation 116, 1128–1136, doi:10.1161/CIRCULATIONAHA.107.710780 (2007).

50 Johansson, M. et al. Cardiac hypertrophy in a dish: a human stem cell based model. Biol Open 9, doi:10.1242/bio.052381 (2020).

51 Mosqueira, D. et al. CRISPR/Cas9 editing in human pluripotent stem cell-cardiomyocytes highlights arrhythmias, hypocontractility, and energy depletion as potential therapeutic targets for hypertrophic cardiomyopathy. Eur Heart J 39, 3879–3892, doi:10.1093/eurheartj/ehy249 (2018).

52 Cheng, Z. et al. Genome-Wide RNAi Screen Identifies Regulators of Cardiomyocyte Necrosis. ACS Pharmacol Transl Sci 2, 361–371, doi:10.1021/acsptsci.9b00052 (2019).

53 Ortiz-Barahona, A., Villar, D., Pescador, N., Amigo, J. & del Peso, L. Genome-wide identification of hypoxia-inducible factor binding sites and target genes by a probabilistic model integrating transcription-profiling data and in silico binding site prediction. Nucleic Acids Res 38, 2332–2345, doi:10.1093/nar/gkp1205 (2010).

54 Morgenstern, C. et al. Biomarkers of NRF2 signalling: Current status and future challenges. Redox Biol 72, 103134, doi:10.1016/j.redox.2024.103134 (2024).

55 Gray, M. O., Karliner, J. S. & Mochly-Rosen, D. A selective epsilon-protein kinase C antagonist inhibits protection of cardiac myocytes from hypoxia-induced cell death. J Biol Chem 272, 30945–30951, doi:10.1074/jbc.272.49.30945 (1997).

56 Pescador, N. et al. Identification of a functional hypoxia-responsive element that regulates the expression of the egl nine homologue 3 (egln3/phd3) gene. Biochem J 390, 189–197, doi:10.1042/BJ20042121 (2005).

57 Higashimura, Y. et al. Up-regulation of glyceraldehyde-3-phosphate dehydrogenase gene expression by HIF-1 activity depending on Sp1 in hypoxic breast cancer cells. Arch Biochem Biophys 509, 1–8, doi:10.1016/j.abb.2011.02.011 (2011).

58 Pescador, N. et al. Hypoxia promotes glycogen accumulation through hypoxia inducible factor (HIF)-mediated induction of glycogen synthase 1. PLoS One 5, e9644, doi:10.1371/journal.pone.0009644 (2010).

59 van Duijvenboden, S. et al. Integration of genetic fine-mapping and multi-omics data reveals candidate effector genes for hypertension. Am J Hum Genet 110, 1718–1734, doi:10.1016/j.ajhg.2023.08.009 (2023).

60 Ramirez, J. et al. Fine mapping of candidate effector genes for heart rate. Hum Genet 143, 1207–1221, doi:10.1007/s00439-024-02684-z (2024).

61 Fong, S. L. & Capra, J. A. Function and Constraint in Enhancer Sequences with Multiple Evolutionary Origins. Genome Biol Evol 14, doi:10.1093/gbe/evac159 (2022).

62 Tejada-Martinez, D. et al. Positive Selection and Enhancer Evolution Shaped Lifespan and Body Mass in Great Apes. Mol Biol Evol 39, doi:10.1093/molbev/msab369 (2022).

63 Chin, I. M., Gardell, Z. A. & Corces, M. R. Decoding polygenic diseases: advances in noncoding variant prioritization and validation. Trends Cell Biol 34, 465–483, doi:10.1016/j.tcb.2024.03.005 (2024).

64 Rao, S., Yao, Y. & Bauer, D. E. Editing GWAS: experimental approaches to dissect and exploit disease-associated genetic variation. Genome Med 13, 41, doi:10.1186/s13073-021-00857-3 (2021).

65 Fulco, C. P. et al. Activity-by-contact model of enhancer-promoter regulation from thousands of CRISPR perturbations. Nat Genet 51, 1664–1669, doi:10.1038/s41588-019-0538-0 (2019).

66 Yao, D. et al. Multicenter integrated analysis of noncoding CRISPRi screens. Nat Methods 21, 723–734, doi:10.1038/s41592-024-02216-7 (2024).

67 Chen, P. B. et al. Systematic discovery and functional dissection of enhancers needed for cancer cell fitness and proliferation. Cell Rep 41, 111630, doi:10.1016/j.celrep.2022.111630 (2022).

68 Murugesan, S. N. & Monteiro, A. Evolution of modular and pleiotropic enhancers. J Exp Zool B Mol Dev Evol 340, 105–115, doi:10.1002/jez.b.23131 (2023).

69 Nord, A. S. et al. Rapid and pervasive changes in genome-wide enhancer usage during mammalian development. Cell 155, 1521–1531, doi:10.1016/j.cell.2013.11.033 (2013).

70 Dickel, D. E. et al. Genome-wide compendium and functional assessment of in vivo heart enhancers. Nat Commun 7, 12923, doi:10.1038/ncomms12923 (2016).

71 Hsiung, C. C. et al. Engineered CRISPR-Cas12a for higher-order combinatorial chromatin perturbations. Nat Biotechnol 43, 369–383, doi:10.1038/s41587-024-02224-0 (2025).

72 Wolfe, J. C., Mikheeva, L. A., Hagras, H. & Zabet, N. R. An explainable artificial intelligence approach for decoding the enhancer histone modifications code and identification of novel enhancers in Drosophila. Genome Biol 22, 308, doi:10.1186/s13059-021-02532-7 (2021).

73 Pradeepa, M. M. et al. Histone H3 globular domain acetylation identifies a new class of enhancers. Nat Genet 48, 681–686, doi:10.1038/ng.3550 (2016).

74 Yan, L. et al. OSAT: a tool for sample-to-batch allocations in genomics experiments. BMC Genomics 13, 689, doi:10.1186/1471-2164-13-689 (2012).

75 Schmidt, D. et al. ChIP-seq: using high-throughput sequencing to discover protein-DNA interactions. Methods 48, 240–248, doi:10.1016/j.ymeth.2009.03.001 (2009).

76 Tohyama, S. et al. Distinct metabolic flow enables large-scale purification of mouse and human pluripotent stem cell-derived cardiomyocytes. Cell Stem Cell 12, 127–137, doi:10.1016/j.stem.2012.09.013 (2013).

77 Zhang, Y. et al. Model-based analysis of ChIP-Seq (MACS). Genome Biol 9, R137, doi:10.1186/gb-2008-9-9-r137 (2008).

78 Kuhn, R. M., Haussler, D. & Kent, W. J. The UCSC genome browser and associated tools. Brief Bioinform 14, 144–161, doi:10.1093/bib/bbs038 (2013).

79 Cunningham, F. et al. Ensembl 2022. Nucleic Acids Res 50, D988–D995, doi:10.1093/nar/gkab1049 (2022).

80 Arnold, M., Raffler, J., Pfeufer, A., Suhre, K. & Kastenmuller, G. SNiPA: an interactive, genetic variant-centered annotation browser. Bioinformatics 31, 1334–1336, doi:10.1093/bioinformatics/btu779 (2015).

81 Labun, K., Montague, T. G., Gagnon, J. A., Thyme, S. B. & Valen, E. CHOPCHOP v2: a web tool for the next generation of CRISPR genome engineering. Nucleic Acids Res 44, W272–276, doi:10.1093/nar/gkw398 (2016).

82 Doench, J. G. et al. Optimized sgRNA design to maximize activity and minimize off-target effects of CRISPR-Cas9. Nat Biotechnol 34, 184–191, doi:10.1038/nbt.3437 (2016).

83 Mandegar, M. A. et al. CRISPR Interference Efficiently Induces Specific and Reversible Gene Silencing in Human iPSCs. Cell Stem Cell 18, 541–553, doi:10.1016/j.stem.2016.01.022 (2016).

84 Burridge, P. W., Holmstrom, A. & Wu, J. C. Chemically Defined Culture and Cardiomyocyte Differentiation of Human Pluripotent Stem Cells. Curr Protoc Hum Genet 87, 21 23 21–21 23 15, doi:10.1002/0471142905.hg2103s87 (2015).

85 Berg, S. et al. ilastik: interactive machine learning for (bio)image analysis. Nat Methods 16, 1226–1232, doi:10.1038/s41592-019-0582-9 (2019).

