## Supplemental Figures for "Mammalian epigenomic conservation of promoters and enhancers in the heart associates with trait-associated variation and impacts cardiomyocyte phenotypes"

---

**SUPPLEMENTAL DATA**

**Figure S1. Basic properties and cross-mapping of promoters, enhancers and primed enhancers identified in heart tissues across 11 mammals** (related to Figure 1) **(page 3)**

**Figure S2. Additional properties of epigenomically-conserved elements and their interplay with trait-associated variants and regulatory signatures** (related to Figure 2) **(p. 4)**

**Figure S3. Selection criteria and genotyping details for targeted enhancers and promoters across three cardiovascular loci** (related to Figure 3) **(p. 6)**

**Figure S4. Differentially-expressed genes and gene ontology associations upon enhancer and promoter deletions for additional time-points during cardiomyocyte differentiation** (related to Figure 4) **(p. 8)**

**Figure S5. Additional details for measurement of hypertrophic and hypoxia-reoxygenation responses in iPSC-cardiomyocytes derived from wild-type and CRISPR knock-out of *CPEB4* and *PRKCE* promoters** (related to Figure 5) **(p. 10)**

**Table S1**: Species and tissue samples used in this study (related to Methods, Figure 1 and Figure S1) **(p. 12)**

**Table S2.** Enrichment analyses for trait-associated genetic variants and regulatory signatures in epigenomically-conserved, primate-specific and human-only regulatory elements (related to Figure 2) **(p.12)**

**Table S3.** Selected epigenomically-conserved regulatory elements harbouring trait-associated variants and overlapping regulatory signatures in cardiomyocytes (related to Figure 3) **(p.12)**

**Table S4.** Differentially expressed genes for CRISPR-KO of selected enhancers and promoters during cardiomyocyte differentiation (related to Figure 4) **(p.12)**

**Table S5.** Gene ontology enrichments for differentially expressed genes across enhancer and promoter perturbations during cardiomyocyte differentiation (related to Figure 4) **(p.12)**

**Table S6.** Guide RNA sequences for CRISPR knock-outs and primers used for genotyping and Sanger sequencing of CRISPR promoter and enhancer deletion clones (related to Figures 3, 4 and 5). (**p.12**)

**Table S7.** Primers used for locus-specific RT-QPCR (related to Figures 3 and 5). (**p. 12**)

### SUPPLEMENTAL FIGURES

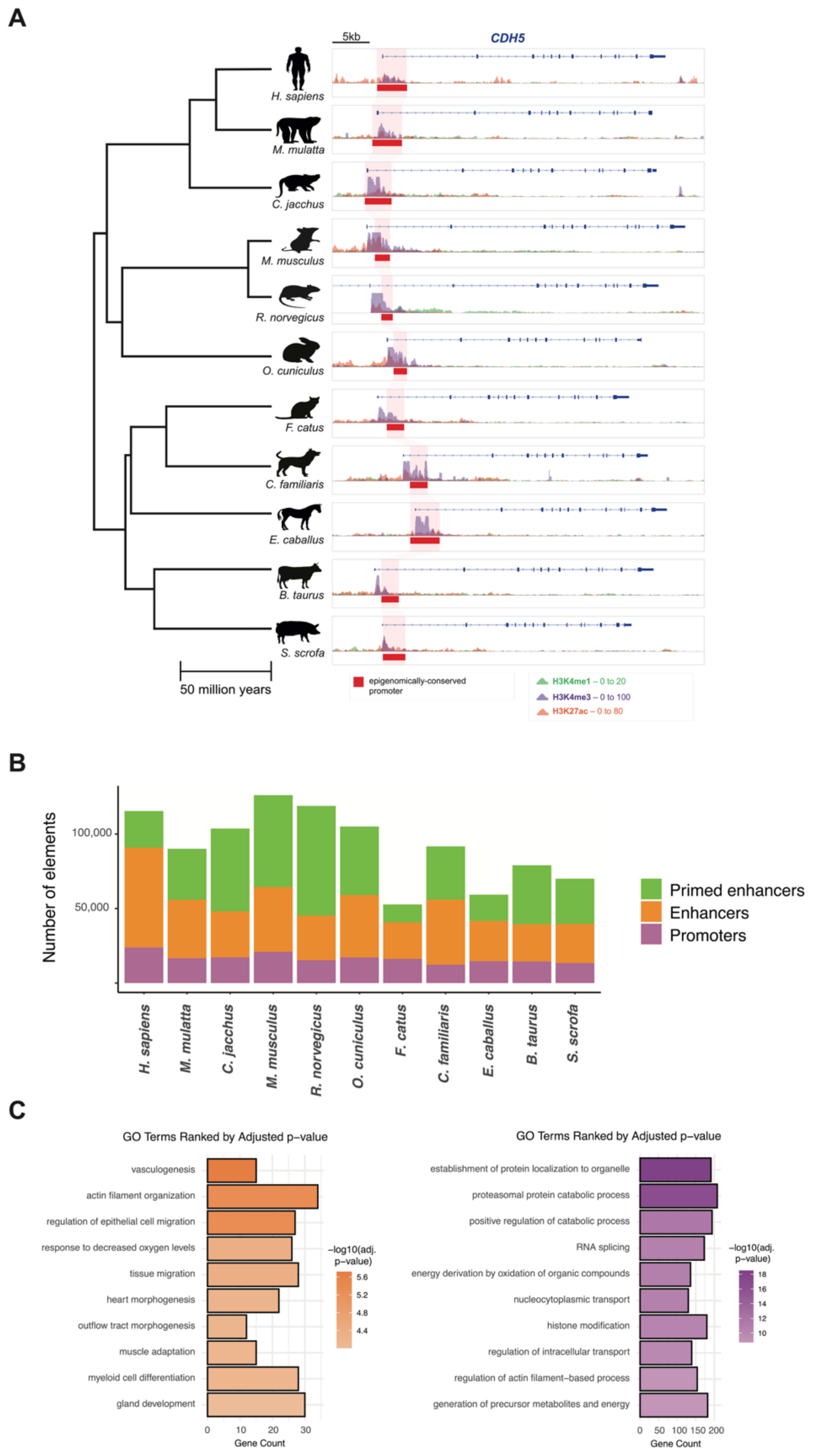


**Figure S1. Basic properties and cross-mapping of promoters, enhancers and primed enhancers identified in heart tissues across 11 mammals** (related to Figure 1)

**A.** Epigenomic signals in heart tissue around the endothelium-expressed gene *Cdh5*, highlighting an epigenomically-conserved promoter region (red bars and shaded area). Details for epigenomic signals and species as in Figure 1A.

**B.** Numbers of promoters (purple), enhancers (orange) and primed enhancers (green) reproducibly identified in each species' heart samples (Methods). [Add average numbers]

**C.** Gene ontology terms associated to epigenomically-conserved enhancers (orange, left) and promoters (purple, right) in the heart. Terms are ranked by statistical significance (FDR-adjusted p-values, Methods).

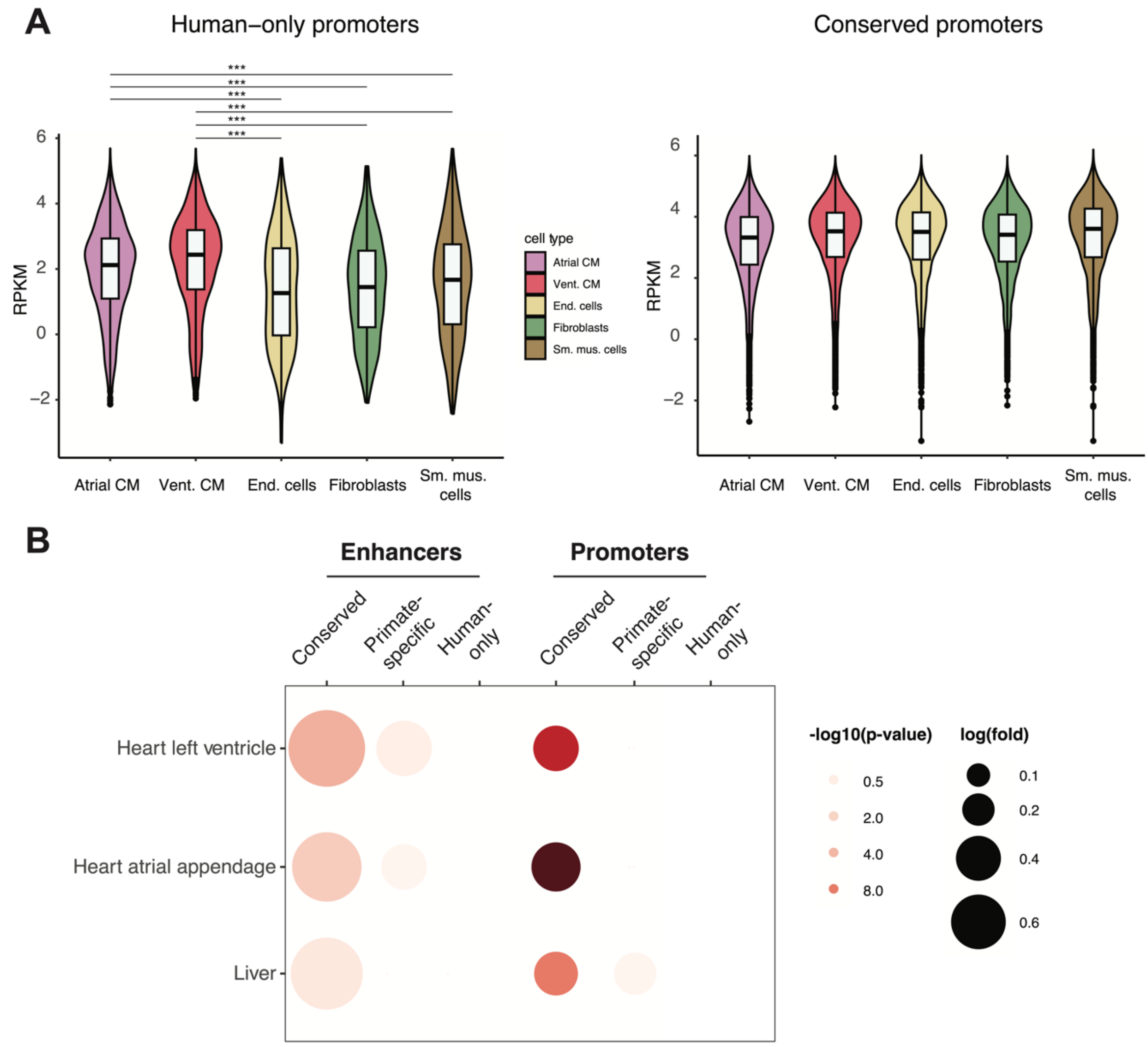


**Figure S2. Additional properties of epigenomically-conserved elements and their interplay with trait-associated variants and regulatory signatures** (related to Figure 2)
**A.** Normalised chromatin accessibility levels across cardiac cell-types for regions overlapping Human-only promoters (left) and Epigenomically-conserved promoters (right).

Shown cardiac cell-types correspond to atrial cardiomyocytes (a_CM), ventricular cardiomyocytes (v_CM), endothelial cells (EC), fibroblasts (FB) and smooth-muscle cells (SM). P-values: Mann Whitney U tests with Benjamini-Hochberg correction for multiple testing, *** < 0.001

**B.** Enrichment of GTEX eQTLs for heart ventricle, heart atrial appendage or liver (y-axis) in epigenomically-conserved or primtate-specific enhancers and promoters (x-axis). P-values: hypergeometric test with Benjamini-Hochberg correction for multiple comparisons.

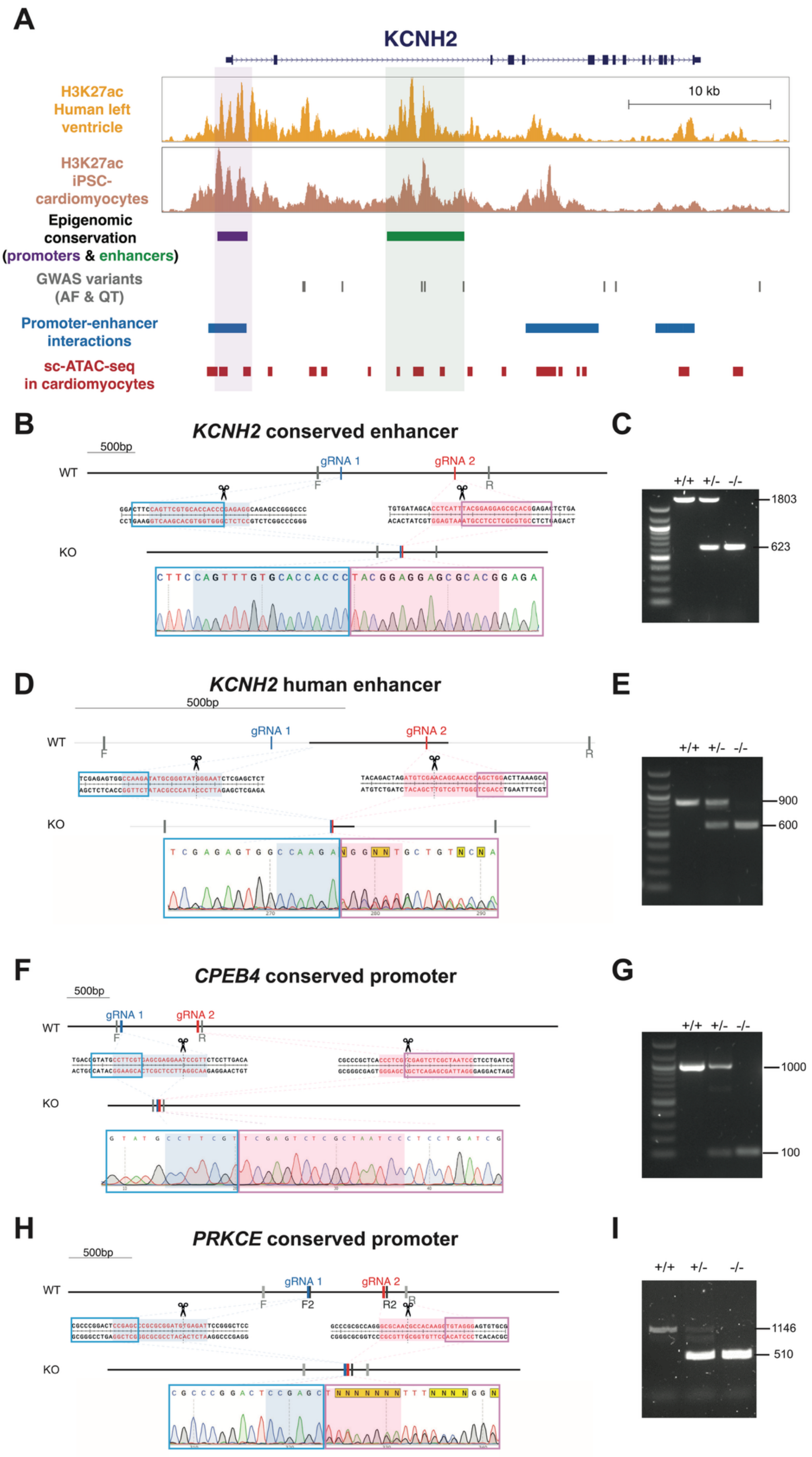
A
KCNH2
H3K27ac
Human left ventricle
10 kb
H3K27ac
iPSC-cardiomyocytes
Epigenomic conservation
(promoters & enhancers)
GWAS variants
(AF & QT)
Promoter-enhancer interactions
sc-ATAC-seq
in cardiomyocytes
B
KCNH2 conserved enhancer
500bp
WT
gRNA 1
gRNA 2
F
R
KO
C
+/+ +/- -/-
1803
623
D
KCNH2 human enhancer
500bp
WT
gRNA 1
gRNA 2
F
R
KO
E
+/+ +/- -/-
900
600
F
CPEB4 conserved promoter
500bp
WT
gRNA 1
gRNA 2
F
R
KO
G
+/+ +/- -/-
1000
100
H
PRKCE conserved promoter
500bp
WT
gRNA 1
gRNA 2
F
F2
R2
R
KO
I
+/+ +/- -/-
1146
510

**Figure S3. Selection criteria and genotyping details for targeted enhancers and promoters across three cardiovascular loci** (related to Figure 3)

**A.** Summary diagram of selection criteria for candidate promoters and enhancers in the heart (Methods), exemplified for the *KCNH2* cardiac locus. Candidate regions were selected based on epigenomic conservation in the heart across mammals, maintenance of H3K27ac enrichments in iPSC-cardiomyocytes and human heart ventricle (top tracks), overlap with cardiovascular GWAS signals and signatures of cardiomyocyte gene regulation in promoter-capture HiC (blue, Promoter-enhancer interactions) and chromatin accessibility (red) datasets.

**B-C.** Representative Sanger sequencing (B) and PCR genotyping (C) of iPSCs harbouring a genomic deletion of a candidate epigenomically-conserved enhancer in *KCNH2*. The left diagram (B) shows guide RNA (gRNA) sequences used for CRISPR/Cas9 genomic deletion relative to the wild-type sequence (top, WT). The bottom chromatogram shows Sanger sequencing results in a knock-out clone. Shaded areas correspond to partial gRNA sequences (blue for gRNA1 and pink for gRNA2). Representative PCR genotyping (C) for wild-type (+/+), heterozygous (+/-) and homozygous (-/-) iPSC clones.

**D-E.** Representative Sanger sequencing (D) and PCR genotyping (E) of iPSCs harbouring a genomic deletion of a control human-only enhancer in *KCNH2*. Labeling as in B-C.

**F-G.** Representative Sanger sequencing (F) and PCR genotyping (G) of iPSCs harbouring a genomic deletion of a candidate epigenomically-conserved promoter in the *CPEB4* locus. Labelling as in B-C.

**H-I.** Representative Sanger sequencing (F) and PCR genotyping (G) of iPSCs harbouring a genomic deletion of a candidate epigenomically-conserved promoter in the *PRKCE* locus. Labelling as in B-C.

See also Table S3 and Table S6.

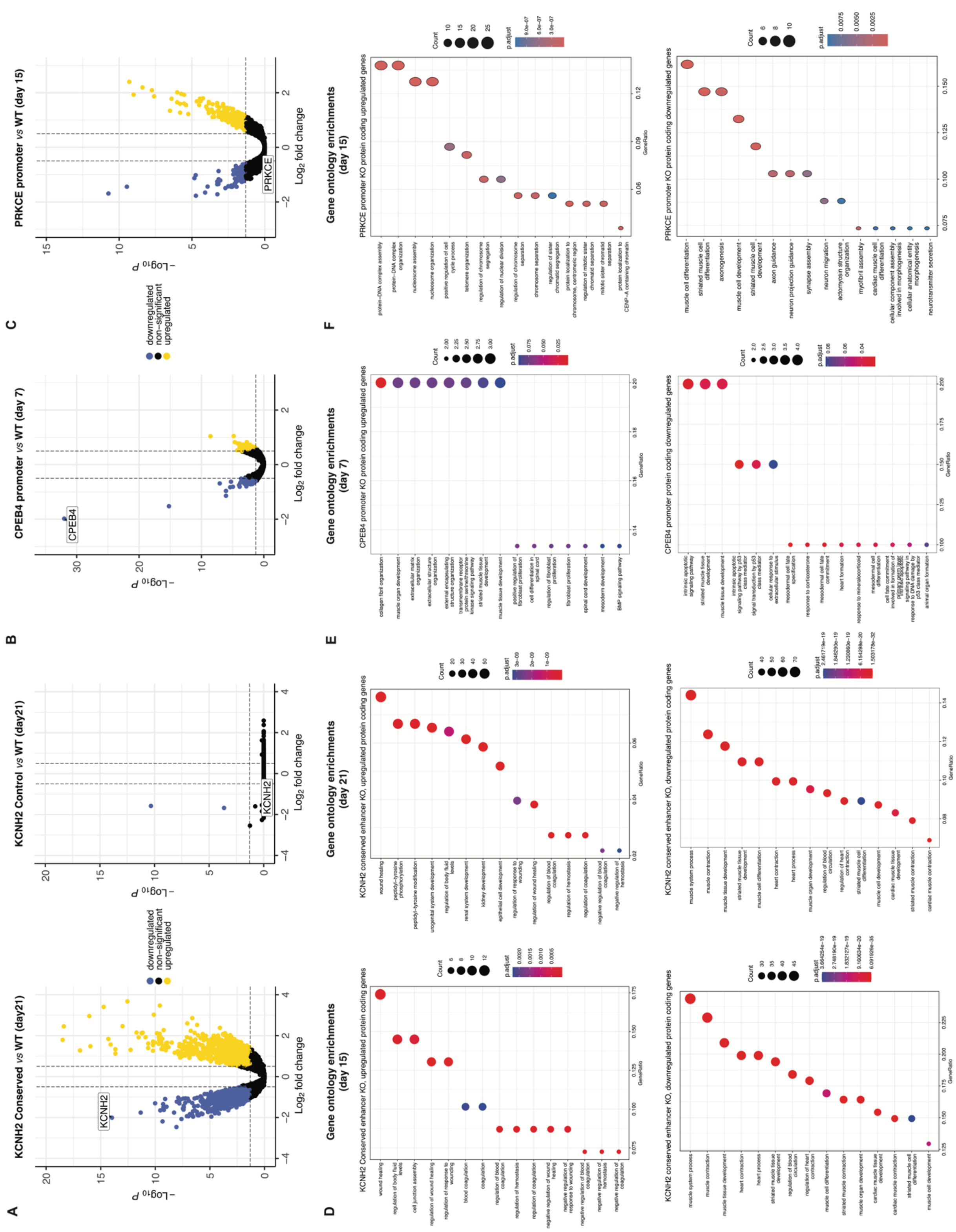


**Figure S4. Differentially-expressed genes and gene ontology associations for additional time-points during cardiomyocyte differentiation for enhancer and promoter deletions** (related to Figure 4)

**A-C.** Differentially expressed genes for additional time-points during cardiomyocyte differentiation of iPSC enhancer and promoter deletions. Volcano plots represent gene expression change (log2 fold changes; x-axis) and statistical significance (-log10(p-value)) for differentially expressed (downregulated, blue; upregulated, yellow; or non-significant, black) and each comparison to wild-type cultures at the indicated time-points, as follows: (A) *KCNH2* conserved enhancer at day 21 (Conserved, left), Kcnh2 human-only enhancer at day 21 (Control, right); (C) *CPEB4* conserved promoter at day 7 (CPEB4 promoter); and (E) *PRKCE* conserved promoter at day 15 (PRKCE promoter).

**D-F**. Gene ontology enrichments for sets of differentially-expressed genes (either upregulated or downregulated) and the indicated comparisons and time-points: (B) Upregulated (top) and downregulated genes (bottom) upon deletion of *KCNH2* conserved (Conserved) at days 15 (left) and 21 (right) of cardiomyocyte differentiation; (D) upregulated (top) and downregulated genes (bottom) upon deletion of *CPEB4* conserved promoter at day 7 of differentiation; and (F) upregulated (top) and downregulated genes (bottom) upon deletion of *PRKCE* promoter at day 15 of differentiation. Gene ontology enrichments for each set are represented as a circle plot (count (size) and adjusted p-value (shade); Methods).

See also Table S5.

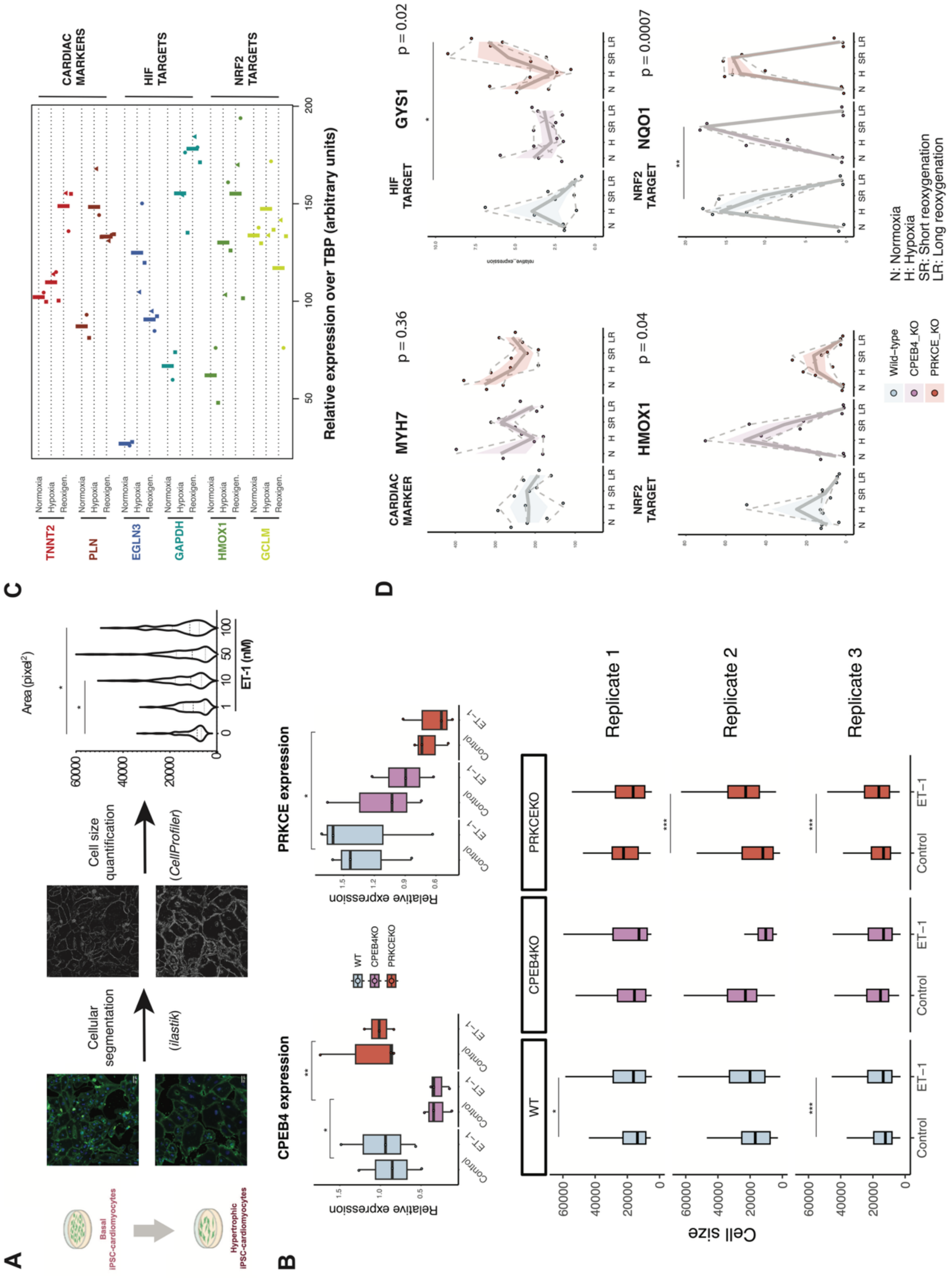


**Figure S5. Additional details for measurement of hypertrophic and hypoxia-reoxygenation responses in iPSC-cardiomyocytes derived from wild-type and CRISPR-KO of *CPEB4* and *PRKCE* promoters**

**A.** Set-up of a high-content microscopy method for quantification of iPSC-derived cardiomyocyte hypertrophy in response to endothelin-1. (Left) Wheat-germ agglutinin and DAPI staining of basal iPSC-cardiomyocytes (top) and cultures treated with 10nM endothelin-1 for 72 hours (bottom). (Center) Cellular segmentation maps obtained with ilastik (Methods) for basal (top) and endothelin-1 treated cultures (bottom). (Right) Cell area distributions for basal cardiomyocytes (Control) and cells treated with the indicated doses of endothelin-1 (ET-1). P-values correspond to sided Mann Whitney U tests with Benjamini-Hochberg correction for multiple comparisons: * < 0.05

**B.** (Top) Real-time quantitative PCR determination of *CPEB4* (left) and *PRKCE* (right) expression levels in cardiomyocytes derived from wild-type (blue), *CPEB4* promoter knock-out (violet) or *PRKCE* promoter knock-out (red) iPSCs. For each genotype, expression levels are shown for basal cultures (Control) and those treated with endothelin-1 (ET-1). P-values across genotypes correspond to 2-way ANOVA with Tukey post-hoc correction: ** < 0.01 , * < 0.05 (Bottom) Distributions of cardiomyocyte cell size across replicate experiments of basal (Control) and hypertrophic (ET-1) cultures derived from iPSCs of wild-type (WT, blue), *CPEB4* promoter knock-out (CPEB4KO, violet) or *PRKCE* promoter knock-out genotype (PRKCEKO, red). P-values for ET-1 response correspond to sided Man Whitney U tests with Benjamini-Hochberg correction for multiple comparisons: *** < 0.001 , * < 0.05

**C.** Set-up of gene expression markers in triplicate iPSC-cardiomyocyte cultures across normoxic conditions (Normoxia, 21% oxygen), 6 hours of 1% oxygen exposure (Hypoxia) and the same followed by 6 hours of reoxygenation in normoxia (Reoxigen.). Relative expression levels are shown in each sample for cardiac (*TNNT2* and *PLN*), hypoxia-responsive (HIF targets *EGLN3* and *GAPDH*) and reoxygenation-responsive genes (NRF2 targets *HMOX1* and *GCLM*). Data from individual replicates is represented as dots, and the average across replicates as thicker vertical bars.

**D.** Gene expression levels for *MYH7* (cardiac marker, top left), *GYS1* (HIF target, top right), *HMOX1* and *NQO1* (both NRF2 targets, bottom) across iPSC-cardiomyocyte samples from the indicated genotypes (wild-type, blue; *CPEB4* promoter knock-out, violet; and *PRKCE* promoter knock-out, red) and treatments (N, normoxia; H, hypoxia; SR: short reoxygenation; and LR: long reoxygenation). P-values: repeated measures two-way ANOVA for interaction between treatment and genotype; Tukey post-hoc correction for pairwise comparisons ** < 0.01 , * < 0.05

### SUPPLEMENTAL TABLES

**Table S1**: Species and genome assemblies used in this study (related to Methods, Figure 1 and Figure S1) **[TableS1.xlsx]**

**Table S2.** Enrichment analyses for trait-associated genetic variants and regulatory signatures in epigenomically-conserved, primate-specific and human-only regulatory elements (related to Figure 2) **[TableS2.xlsx]**

**Table S3.** Selected epigenomically-conserved regulatory elements harbouring trait-associated variants and overlapping regulatory signatures in cardiomyocytes (related to Figure 3) **[TableS3.xlsx]**

**Table S4.** Differentially expressed genes for CRISPR-KO of selected enhancers and promoters during cardiomyocyte differentiation (related to Figure 4) **[TableS4.xlsx]**

**Table S5.** Gene ontology enrichments for differentially expressed genes across enhancer and promoter perturbations during cardiomyocyte differentiation (related to Figure 4) **[TableS5.xlsx]**

**Table S6.** Sequences for guide RNAs, primers used for genotyping and Sanger sequencing primers of CRISPR promoter and enhancer deletion clones (related to Figures 3, 4 and 5). **[TableS6.xlsx]**

**Table S7**. Primers used for locus-specific RT-qPCR (related to Figures 3 and 5). **[TableS7.xlsx]**
